# Evolutionary Remodeling of Ubiquinone Biosynthesis in *Toxoplasma gondii* Reveals an Essential Bi-functional Monooxygenase

**DOI:** 10.64898/2026.02.10.705114

**Authors:** Baihetiya Baierna, Taufiq Rahman, Scott Latimer, Gilles J Basset, Silvia NJ Moreno

## Abstract

Apicomplexan parasites like *Toxoplasma gondii* harbor a highly divergent mitochondrial proteome, much of which remains uncharacterized despite its essentiality for parasite survival. One such critical pathway is ubiquinone (UQ) biosynthesis. Here, we characterize the UQ synthesis machinery in *T. gondii* and show that conserved enzymes, TgCoq3 and TgCoq5, are essential for growth and mitochondrial function, forming a multi-protein complex. Using proximity labeling and subcellular fractionation, a strategy suited for detecting proteins of low abundance, we identify TgCoqFAD, a unique FAD-dependent monooxygenase required for UQ synthesis. Unlike canonical eukaryotic systems that employ multiple monooxygenases to modify specific carbons on the UQ aromatic ring, TgCoqFAD catalyzes two distinct hydroxylation steps, an activity not previously reported in eukaryotes. Molecular docking and chemical screening identified TgCoqFAD inhibitors that impair tachyzoite growth and bradyzoite viability. These findings reveal a streamlined and divergent UQ biosynthesis pathway in apicomplexans and establish TgCoqFAD as a promising antiparasitic target.

## Introduction

*Toxoplasma gondii* is an obligate intracellular parasite of the phylum Apicomplexa, which also includes major human pathogens such as *Plasmodium* and *Cryptosporidium spp. T. gondii* infects nearly one-third of the global population^1^. Following initial infection, *T. gondii* progresses through an acute stage characterized by rapid replication of tachyzoites, which can disseminate systemically and cause tissue damage. This is followed by a chronic stage, during which the parasite persists as latent bradyzoite in tissue cysts primarily in muscle and neural tissues^2^. Reactivation of latent infection in immunocompromised individuals can result in severe tissue damage in the heart^3^, brain^4^, and lungs^5^. Current treatment for toxoplasmosis, such as pyrimethamine combined with sulfadiazine^6^, are limited by significant toxicity and frequent relapses upon cessation^6–8^. Consequently, there is a pressing need for novel therapeutics and parasite-specific targets essential for both acute and chronic stages of infection.

The mitochondrion of *T. gondii* represents a compelling therapeutic target due to its central role in parasite viability and the presence of essential enzymes that are highly divergent from those of the host^9^. Notably, atovaquone, a drug used to treat malaria, toxoplasmosis and babesiosis, targets the mitochondrial electron transport chain (ETC) at the ubiquinone (UQ): cytochrome c reductase (complex III)^10,11,12^. This underscores the potential of the UQ biosynthesis as a promising target for therapeutic intervention.

Ubiquinone (UQ), also known as coenzyme Q, is a redox-active lipid composed of a water soluble quinone head derived from *para*-hydroxybenzoate (PHBA) and a polyisoprene tail that anchors it to the inner mitochondrial membrane^13,14^. In the *T. gondii* ETC, UQ functions as the central electron carrier linking multiple dehydrogenases to complex III. In this context, because the parasite lacks a canonical complex I, electron input into the UQ pool instead occurs through succinate dehydrogenase (complex II), and additional enzymes, including malate-quinone oxidoreductase (MQO), dihydroorotate dehydrogenase (DHODH), type II NADH dehydrogenase and glycerol-3-phosphate dehydrogenase (G3PDH)^15–18^. Beyond its role in oxidative phosphorylation, UQ is also required for pyrimidine synthesis, fatty acid β-oxidation, and redox homeostasis^19–24^.

Most of our understanding of UQ biosynthesis derives from studies in yeast, mammals, and bacteria^25^, where a largely conserved pathway involving a series of enzymes has been defined^26^ (**Supplementary Fig. 1**). In eukaryotes, the first two enzymes of this pathway, Coq1, a soluble matrix protein, and Coq2 (COQ1 and COQ2 in mammals), an inner membrane bound protein, synthesize the isoprene tail and attach it to the benzoquinone ring, respectively^26^. Subsequent ring modifications include decarboxylation, followed by a series of hydroxylation and methylation steps to ultimately yield the fully substituted UQ molecule (**Fig. 1**). The hydroxylation of the benzoquinone ring differs between organisms. In yeast and mammals, C5 hydroxylation is carried out by an FAD-dependent monooxygenases (Coq6/COQ6 respectively), while C6 hydroxylation is catalyzed by a diiron hydroxylase (Coq7/COQ7 respectively)^25^. In contrast, bacteria and plants use FAD-dependent monooxygenases to perform hydroxylation at different carbon positions^25,27^. In yeast, the enzymes responsible for these ring decorations, Coq3-Coq9 and Coq11, assemble into a large multi-protein complex known as the UQ synthome^26^. A similar protein complex is also present in mammals (the UQ complex)^28^, and in bacteria (the UQ metabolome)^29^. These complexes are thought to enhance the catalytic efficiency and sequester potentially toxic intermediates, preventing oxidative damage. Despite the central role of UQ in cellular metabolism, the mechanism of UQ ring modifications and the presence and organization of a UQ synthesis complex have not been investigated in *T. gondii* or other apicomplexan parasites to date.

**Figure 1.**
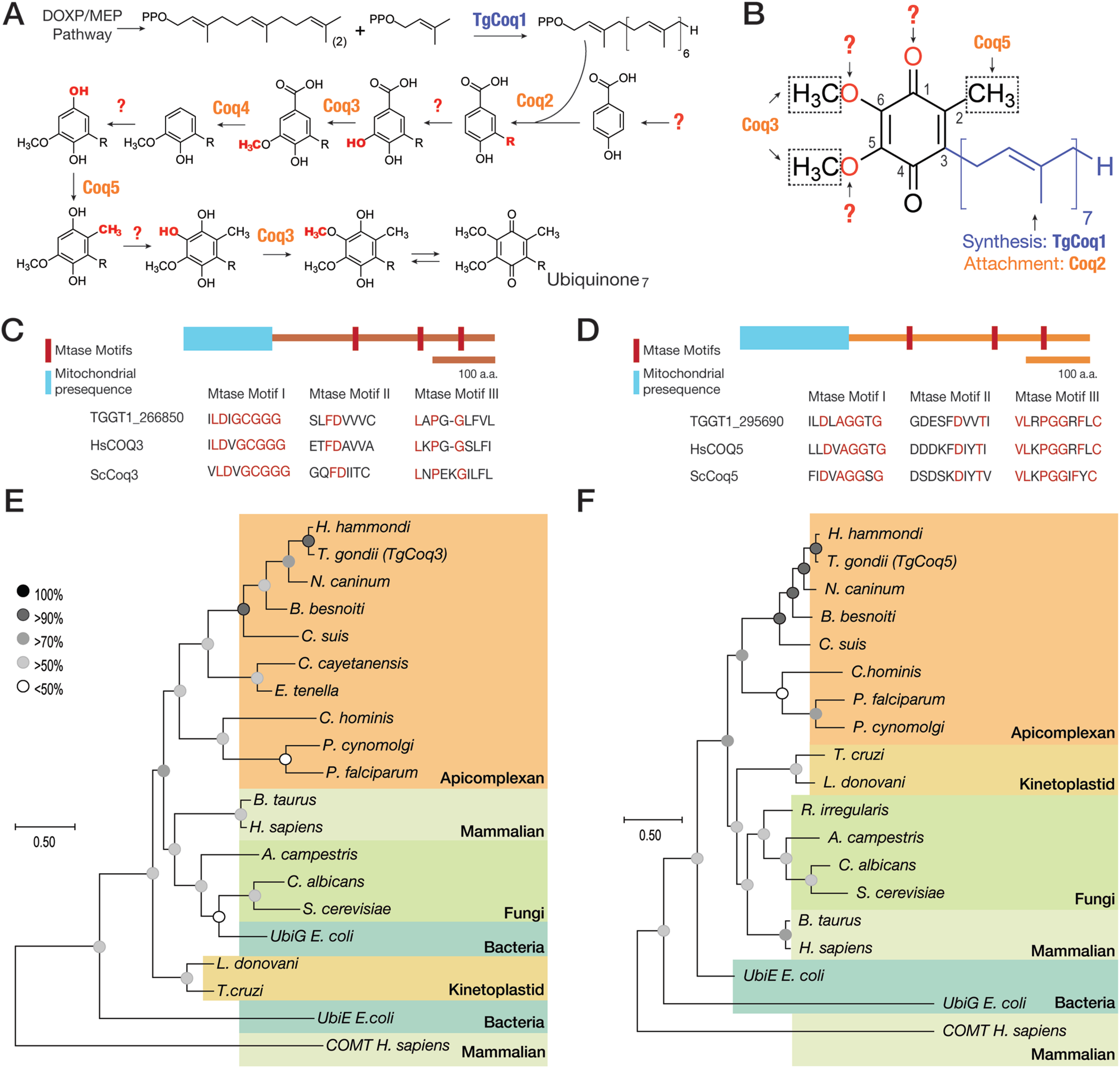
Predicted ubiquinone biosynthesis pathway and evolutionary analysis of Coq enzymes in *T. gondii*. (A) Schematics of the predicted ubiquinone synthesis pathway in *T. gondii* adapted from the yeast pathway^26^. Enzymes homologous to those of the yeast pathway are shown in orange, and steps for which no gene has been predicted are indicated by question marks. (B) Structure of UQ and the enzymes involved in the ring group modification that have homologs in *T. gondii* (orange). Question mark indicates unidentified genes. (C) Predicted domain arrangement of the TgCoq3 (TGGT1_266850) protein with the mitochondrial signal peptide indicated in blue and the methyltransferase motifs (Mtase) in red. Conserved methyltransferase motif sequences^94^ were compared with the *H. sapiens* and *S. cerevisiae* Coq3 protein motifs. (D) Predicted domain arrangement of the TgCoq5 (TGGT1_295690) protein with the predicted mitochondrial signal peptide indicated in blue and methyltransferase motifs (Mtase) in red. Conserved methyltransferase motif sequences^94^ were compared with *H. sapiens* and *S. cerevisiae* Coq5 protein motifs. (E) Phylogenetics of Coq3 in different organisms. UbiE (bacterial homolog of Coq5) and COMT (Catechol-*O*-methyltransferase) are used as outgroups. Sequences used are listed in **Supplementary table 2**. (F) Phylogenetics of Coq5 in different organisms. UbiG (bacterial homolog of Coq3) and COMT are used as outgroups. Sequences used are listed in **Supplementary table 2**.

Our previous work demonstrated that TgCoq1 can be targeted by lipophilic bisphosphonates^30^, commonly used to treat osteoporosis^31^, supporting the druggability of this pathway. However, bioinformatic analyses have failed to identify all the enzymes required for UQ ring modifications in *T. gondii*, suggesting that the parasite may use novel and divergent enzymes for these steps. Given the divergence of other mitochondrial protein complexes in *T. gondii* and the presence of parasite-specific subunits identified within each ETC complex^32,33^, we hypothesized that UQ synthesis in apicomplexans also involves unique enzymes assembled into a stable protein complex.

In this study we present the first biochemical characterization of the UQ biosynthesis machinery in *T. gondii*. We investigate the potential assembly of these enzymes into a multi-protein complex, with a focus on identifying parasite-specific enzymes that differ from those found in mammals and yeast. Our work aims to expand the understanding of mitochondrial metabolism in apicomplexan parasites and to explore novel targets for therapeutic intervention.

## Results

### The ubiquinone ring modifying enzymes of *T. gondii*

To identify potential enzymes involved in the UQ synthesis (**Fig. 1A**), we first used the *Saccharomyces cerevisiae* UQ synthesis protein sequences and performed a BLAST^34^ and HHPRED^35^ search of the *T. gondii* proteome^36^. For some of the corresponding proteins, strong homologs were found in *T. gondii* (**Supplementary Table 1).** Our laboratory had previously characterized TgCoq1 and found that it is responsible for the synthesis of the isoprenoid unit, heptaprenyl diphosphate^30^. However, bioinformatic searches could not identify obvious homologs for Coq6, Coq7 or Coq9 (**Fig. 1A and Supplementary Table 1**) which carry out essential enzymatic steps of the synthesis of UQ, or Coq11 (**Fig. 1B and Supplementary Fig. 1**). Interestingly, we identified homologs of Yah1 and Arh1, which provide electrons to Coq6 in *S. cerevisiae*^26^. Based on this, we hypothesized that *T. gondii* possesses a monooxygenase capable of catalyzing the required hydroxylation steps (**Fig. 1B**), but that it is likely highly divergent from its fungal and mammalian counterparts.

To characterize UQ synthesis, we focused on two genes *TGGT1_266850* and *TGGT1_295690*, annotated as 3-demethylubiquinone-9 3-*O*-methyltransferase (*TgCoq3*), and ubiquinone/menaquinone biosynthesis methyltransferase subfamily protein (*TgCoq5*) respectively (**Supplementary Table 1**). These enzymes catalyze key methylation steps in the UQ biosynthesis pathway: Coq3 functions as an *O*-methyltransferase, acting at the C5 and C6 positions of the aromatic ring, while Coq5 serves as a *C*-methyltransferase (**Fig. 1B**). Sequence alignment with yeast (Coq3 and Coq5) and human (COQ3 and COQ5) homologs revealed low identity (TgCoq3: 37.31% human, 27.71% yeast; TgCoq5: 40% human, 32.51% yeast). Despite this divergence, both protein sequences retain conserved methyltransferase motifs (**Fig. 1C** and **Fig. 1D**), supporting their predicted enzymatic function. Phylogenetic analyses including representative animal, plant, yeast, and apicomplexan homologs showed that TgCoq3 and TgCoq5 cluster with UQ synthesis O- and C-methyltransferases, respectively, and are most closely related to other apicomplexan proteins predicted to function in UQ synthesis (**Fig. 1E and F**). Both enzymes remain distant from mammalian homologs, highlighting their evolutionary divergence.

### TgCoq3 and TgCoq5 are important for *T. gondii* growth and UQ synthesis

Subcellular localization of TgCoq3 (TGGT1_266850) and TgCoq5 (TGGT1_295690) was initially predicted to be targeted to the mitochondrion by LOPIT (Localization of Organelle Proteins by Isotope Tagging) analysis^37^ (**Supplementary Table 1**). To validate these predictions, we tagged the C-terminus of both genes with a triple HA tag and isolated clonal lines expressing *TgCoq3-HA* and *TgCoq5-HA*. Immunofluorescence microscopy analysis (IFA) of these lines showed partial co-localization of both proteins with the mitochondrial marker TOM40^38^ (**Supplementary Fig. 2A**). However, because the signal was relatively weak, the localization was not fully convincing. To improve detection, we generated polyclonal antibodies against both proteins. For this, we cloned both *TgCoq3* and *TgCoq5* genes in bacterial expression plasmids and purified the respective recombinant proteins and used them to immunize animals. We generated and purified αTgCoq3 in mouse and αTgCoq5 in guinea pigs which were used to confirm the mitochondrial localization of both proteins **(Supplementary Fig. 3**).

To study the role of TgCoq3 and TgCoq5 in *T. gondii*, we generated conditional knockdown lines by inserting a tetracycline (ATc) response element at the promoter region^39^ of each gene, followed by pyrimethamine selection and clonal isolation (*iΔTgCoq3* and *iΔTgCoq5*) (**Fig. 2A**). Expression of both proteins was reduced 24 h after the addition of ATc to the cultures and completely ablated by 48 h as shown by western blot analysis and IFAs (**Supplementary Fig. 2B-C and Fig. 2B**). We next studied the role of both proteins in the *T. gondii* lytic cycle with plaque assays, in which the parasite engages in repetitive lytic cycles resulting in host cell monolayer lysis forming plaques that can be visualized by staining with crystal violet. Both mutants, *iΔTgCoq3* and *iΔTgCoq5* formed significantly smaller plaques when grown in the presence of ATc (**Fig. 2C-2D**). To facilitate measurement of replication and growth over several days, we expressed a cytosolic red fluorescent protein (tdTOMATO) in both mutants (*iΔTgCoq3-RFP* and *iΔTgCoq5-RFP*). This allowed us to quantify growth by measuring red fluorescence over time. Parasite numbers were significantly reduced in the *iΔTgCoq3-RFP* and *iΔTgCoq5-RFP* mutants when grown in the presence of ATc (**Figs. 2E** and **2F**, *+ATc*). In a separate replication assay, ATc treatment of the mutants for three days resulted in a significant replication defect (**Supple**mentary Fig. 2D).

**Figure 2.**
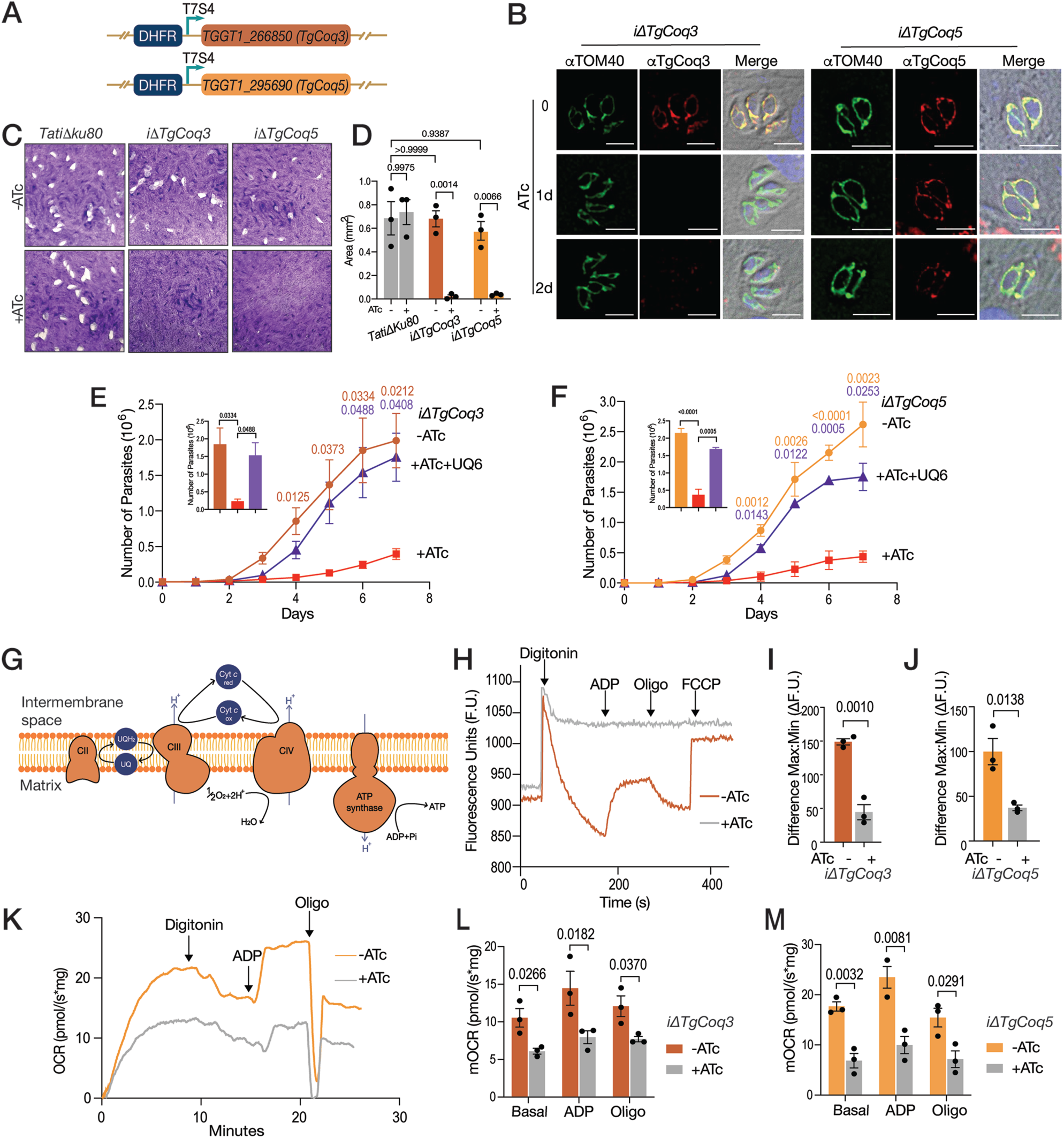
TgCoq3 and TgCoq5 are essential for the lytic cycle, mitochondrial function and UQ synthesis. (A) Schematics showing the genetic modification of the conditional mutants. The 5’ cassette includes a DHFR drug resistant marker, a tetracycline response element and a SAG4 promoter. The primers used are listed in **Supplementary Table 7**. (B) IFAs of *iΔTgCoq3* and *iΔTgCoq5* cell lines. Parasites were incubated with ATc for 0, 1, or 2 days. αTOM40 was used as mitochondrial marker, and αTgCoq3 or αTgCoq5 were used to visualize TgCoq3 or TgCoq5 respectively. Bars are 5 µm. (C) Representative plaques of the *iΔTgCoq3* and *iΔTgCoq5* lines cultured in the presence or not of ATc. Plates were fixed and stained at day 7. (D) Quantification of plaque areas from three biological replicates. Statistical significance was evaluated by two-way ANOVA. Each point represents a biological replicate, and error bars indicate SEM. (E) Growth assay of *iΔTgCoq3-RFP* <u>-ATc</u>: no ATc; <u>+ATc</u>: parasites pretreated with ATc for 3 days before infection and cultured with ATc; <u>+ATc+UQ_6_</u>: same as +ATc, plus 10 µM UQ_6_ added to the medium. One-way ANOVA analysis was used, n=3. Data is presented as mean ± SEM. (F) Growth assay with the *iΔTgCoq5* mutant. Conditions were like the ones in E. One-way ANOVA analysis from three biological replicates, n=3. (G) Scheme of the *T. gondii* Electron Transport Chain. (H) Representative trace of membrane potential measurements in *iΔTgCoq3* parasites using 2.5 µM safranin O, with 1 mM succinate present in the buffer as the respiratory substrate. <u>-ATc</u>: no ATc (*brown trace*); <u>+ATc</u>: parasites cultured with ATc for 2 days (*grey trace*). <u>Oligo</u>: 1 µg/mL oligomycin. (I) *iΔTgCoq3* mitochondrial membrane potential quantification of the differences between the maximum fluorescence (after depolarization with 5 µM FCCP) and the minimum fluorescence (after 30 µM digitonin permeabilization). Student’s t test was used, n=3. Each point represents a biological replicate, and the error bars represent SEM. (J) *iΔTgCoq5* mitochondrial membrane potential quantification, <u>+ATc</u>: ATc for 3 days, other conditions are the same as the ones used in I. (K) Representative tracing of the oxygen consumption rate of the *iΔTgCoq5* mutant, -<u>ATc</u>: no ATc (*orange trace*); <u>+ATc</u>: parasites cultured with ATc for 2 days (*grey trace*). (L) *iΔTgCoq3* mOCR quantification. Basal, ADP-stimulated (10 µM), and oligomycin-inhibited mOCR were quantified from three biological replicates. Two-way ANOVA was used for statistical analysis. Each point represents a biological replicate, and bars indicate SEM. (M) mOCR was quantified from three biological replicates. Two-way ANOVA was used for statistical analysis. Each point represents a biological replicate, and error bars indicate SEM. For **D**-**F** and **I**, **J**, **L**, and **M**, significance was considered when *p*-value is less than 0.05.

To validate the function of TgCoq3 and TgCoq5 in UQ synthesis, we added ATc to the cultures to knockdown the respective genes and supplemented the medium with 10 µM UQ_6_, the end product of the yeast UQ synthesis pathway (**Fig. 2E** and **Fig. 2F**, *+ATc+UQ_6_*). Importantly, the growth repression of both mutants by ATc, was rescued by UQ_6_, supporting the role of TgCoq3 and TgCoq5 in UQ synthesis. However, the recovery was not 100%, possibly due to the limited permeability of UQ_6_ or a preference of *T. gondii* for UQ_7_ in mediating its cellular functions.

### TgCoq3 and TgCoq5 are required for mitochondrial function

UQ transfers electrons from complex II to cytochrome c through complex III in the inner mitochondrial membrane (**Fig. 2G**). This electron transfer is linked to proton transport from the matrix to the intermembrane space, generating a membrane potential. We used safranin *O*^30,40^, a fluorescent dye that inserts into energized mitochondrial membranes, to measure membrane potential. Permeabilizing parasites in suspension with digitonin allowed safranin to enter the cells. In the presence of succinate, the mitochondrial membrane becomes energized, leading to accumulation of safranin and a corresponding decrease in fluorescence (**Fig. 2H**, *brown trace*). Addition of ADP to stimulate ATP synthesis resulted in depolarization of the membrane, which is observed as an increase in fluorescence. Inhibition of ATP synthase by oligomycin restored the membrane potential, observed as a decrease in fluorescence. Lastly, addition of Carbonyl cyanide-p-trifluoromethoxy phenylhydrazone (FCCP) dissipated the membrane potential (Fig**. 2H**, *brown trace*). We next measured the membrane potential of the *iΔTgCoq3* (*+ATc*) mutant, which showed to be significantly unresponsive to all additions (**Fig. 2H** and **Fig. 2I**, *gray trace and bar*). Analysis of the membrane potential of *iΔTgCoq5* (*+ATc*) revealed a similar phenotype (**Fig. 2J**, *gray bar*).

The ETC uses oxygen as its terminal electron acceptor, so we next assessed oxygen consumption of the mutants using an Oroboros O2K respirometer, which measures the rate of mitochondrial oxygen consumption (mOCR). Permeabilizing the plasma membrane with digitonin allowed for the substrate succinate to access and energize the mitochondria, enabling the measurement of basal mOCR (**Fig. 2K**, *orange trace*). Addition of ADP increased the mOCR as it stimulated oxidative phosphorylation. Lastly, inhibition of ATP synthesis with oligomycin reduced the rate of oxygen consumption to a minimal mOCR. All of the characteristics of healthy mitochondrial oxygen consumption were defective in the *iΔTgCoq3* and *iΔTgCoq5* mutants when grown in the presence of ATc (**Fig. 2K-M**, *gray trace and bars*). Taken together, both TgCoq3 and TgCoq5 are essential for *T. gondii* mitochondrial functions.

### The presence of an UQ synthesis complex in *T. gondii*

To investigate the organization and potential interaction between the UQ synthesis enzymes (Fig. 3A), we performed Blue Native PAGE (BN PAGE) analysis. BN PAGE was used for the detection of mitochondrial membrane complexes, and this method has been used to visualize ETC complexes in *T. gondii*^32,41^. Lysates from *TgCoq3-3HA* and parental strain prepared under native conditions, were separated by native PAGE, transferred to a membrane, and probed with αHA antibody, which showed no signal. However, protein transfer was confirmed by probing the membrane with the F1β antibody, which recognizes a subunit of the ATP synthase, revealing a clear reaction corresponding to Complex V (**Supplementary Fig. 4A**). Considering that both TgCoq3 and TgCoq5 are expressed at very low levels, we thought that this could explain the absence of detectable signals. To overcome this issue, we next enriched mitochondrial membranes through subcellular fractionation, using a differential centrifugation protocol that separates organelles based on their density (**Supplementary Fig. 4B**). To identify enriched mitochondrial fractions, we measured succinate cytochrome *c* reductase activity, a mitochondrial marker (**Supplementary Fig. 4C-D**). The P2 fraction showed the highest enrichment of activity, compared to the S1 (total lysate) fraction (**Supplementar**y **Fig. 4C**). We then performed BNPAGE using the P2 fraction, however, the signal was still undetectable. We considered the possibility that the epitopes of TgCoq3 and TgCoq5 could be hidden within the hypothetical complex of *T. gondii*. To address this issue, protein complexes separated by BN-PAGE were subjected to second-dimension SDS-PAGE to resolve individual subunits. Under these conditions, both TgCoq3 and TgCoq5 were detected as components of a high molecular weight (>1,000 kDa) complex (**Fig. 3B**).

**Figure 3.**
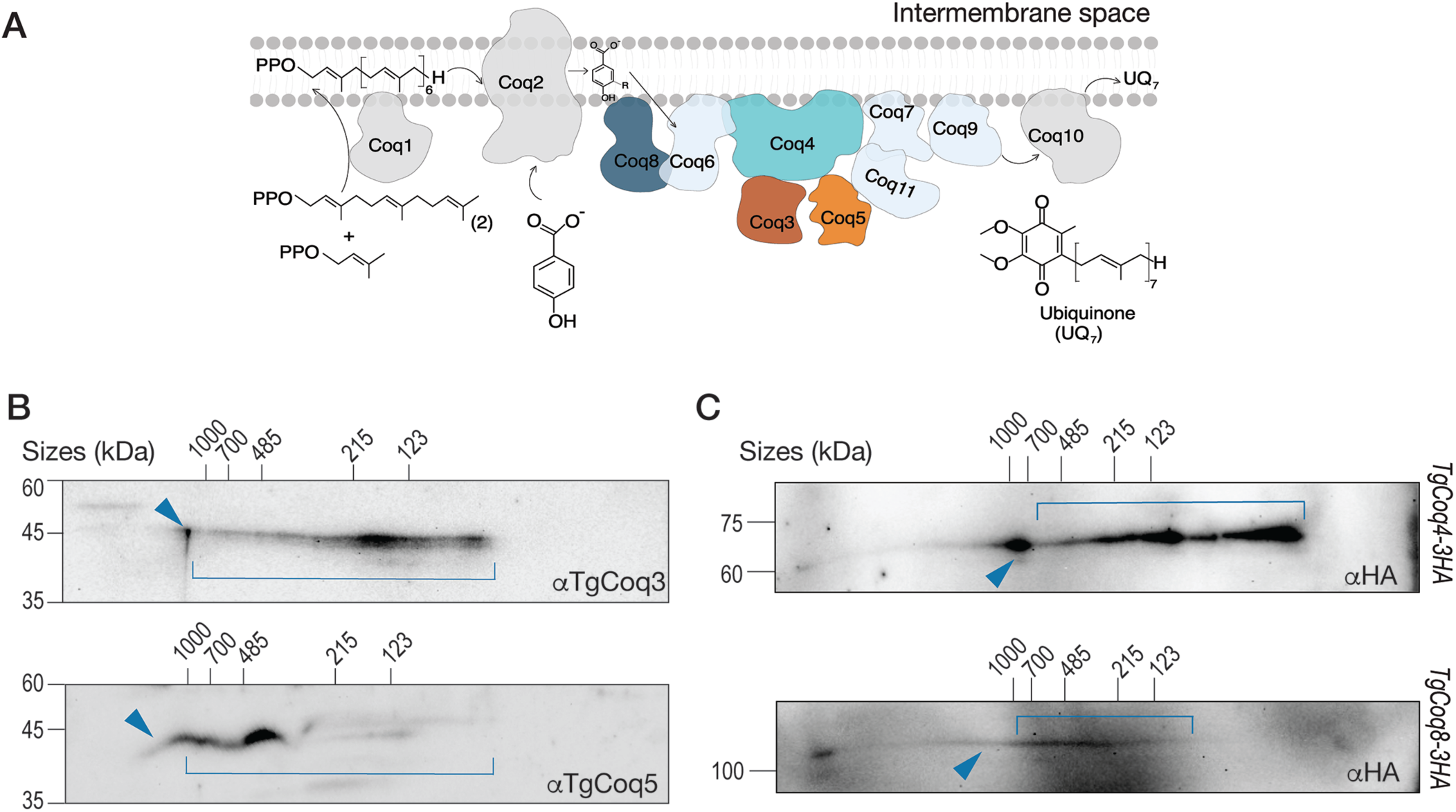
Ubiquinone (UQ) synthesis enzymes in *T. gondii* form a high molecular weight complex. (A) Schematic of the putative UQ synthesis complex. Proteins with homologs in *T. gondii* are shown in solid colors, while proteins lacking identifiable homologs are shown in light blue. (B) 2D BN/SDS-PAGE analysis (details in the methods section) using 100 μg of solubilized P2 fraction obtained using the protocol shown in Supplementary Fig. 4B. Membranes probed with αTgCoq3 and αTgCoq5 antibody. (C) 2D BN/SDS-PAGE analysis using *TgCoq4-3HA* and *TgCoq8-3HA* lysates (protocol detailed in the method section). Membranes probed with αHA antibody. Blue arrowhead indicates the UQ synthesis complex, and blue brackets indicates complex or disassembly intermediates.

To further validate the presence of a UQ complex and to test physical interactions among its components, we focused on TgCoq4 (TGGT1_275980) and TgCoq8 (TGGT1_203940), two conserved Coq proteins predicted to function as structural and regulatory elements of the UQ biosynthetic machinery. We generated *TgCoq4-HA* and *TgCoq8-3HA* lines (**Supplementary Fig.5A and 5B**) and confirmed their mitochondrial localization by IFA (**Supplementary Fig. 5C and 5D**). Reciprocal co-IP experiments revealed interactions between TgCoq4, TgCoq5, and TgCoq8 (**Supplementary Fig. 5E and 5F**), supporting the existence of a multi-protein UQ synthesis complex in *T. gondii*.

To further characterize the organization of the complex, we performed 2D BN/SDS-PAGE using lysates from *TgCoq4-3HA* and *TgCoq8-3HA* (**Fig. 3C**). Contrary to TgCoq3, which could only be detected using the P2 fraction, both TgCoq4 and TgCoq8 were detected in a protein complex larger than 1 MDa using whole cell lysates, likely due to their higher abundance or epitope orientation. However, each protein displayed different lower molecular weights species (indicated with blue brackets), which may indicate different assembly intermediates or partial disassembly during sample preparation. Together, these data provide the first evidence that *T. gondii* contains a multi-subunit UQ synthesis protein complex.

### Novel enzymes of the UQ synthesis pathway identified by TurboID pulldown and subcellular fractionation

The proteome database failed to reveal several enzymes involved in the UQ synthesis pathway, including counterparts to key yeast enzymes responsible for catalyzing essential steps. We hypothesize that these enzymes are present in *T. gondii* but are highly divergent making sequence analyses unrewarding. We initially performed αHA immunoprecipitation assays using whole cell lysates from *TgCoq3-3HA* and *TgCoq5-3HA*, however, the results were unproductive, likely due to the low expression levels of these proteins. Since we confirmed that TgCoq3 and TgCoq5 are both part of a large protein complex, we tagged their C-terminus with a biotin ligase, TurboID^42^, and isolated the clonal mutants *TgCoq3-TID* and *TgCoq5-TID* (**Fig. 4A**).

**Figure 4.**
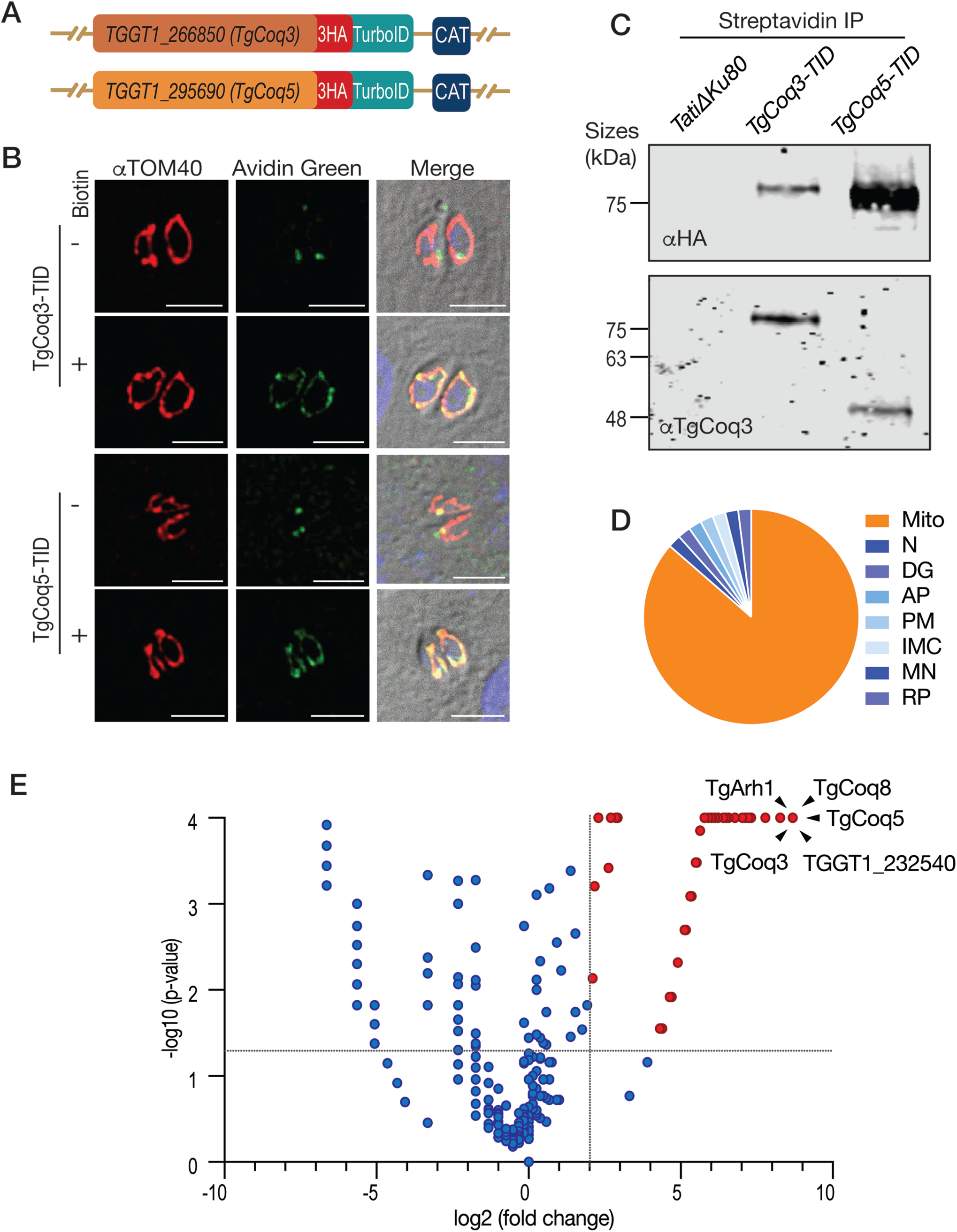
Identification of candidate UQ synthesis enzymes by TurboID and subcellular fractionation. (A) Scheme showing the tagging of TgCoq3 and TgCoq5 with TurboID. CAT: chloramphenicol Acetyl Transferase gene for selection. (B) IFA of tagged cell lines incubated with DMSO or 50 μM biotin for two hours. Biotinylated proteins were probed with avidin green and TOM40 was used as mitochondrial marker. Scale bars are 5 µm. (C) Western blot analysis of eluates from streptavidin co-immunoprecipitation of *TatiΔku80*, *TgCoq3-TID*, and *TgCoq5-TID* parasites incubated with biotin for 2 h. Blots were probed with αHA (top) and αTgCoq3 (bottom). (D) Pie chart showing the predicted localization by LOPIT^39^ of the significantly (*p*<0.05) enriched hits with a fold-change (TgCoq5-TID/FbxO14-TID) higher than or equal to 2. Mito: mitochondria, N: nucleus, DG: dense granules, AP: apicoplast, PM: plasma membrane, IMC: inner membrane complex, MN: microneme, RP: rhoptry. (E) Volcano plots of the peptides detected in the eluates graphed based on their fold-change (TgCoq5-TID/FbxO14-TID) and their *p*-values. Red dots indicate peptides significantly enriched (at least 2-fold) in the eluate of *TgCoq5-TID*. The black arrowhead points at TgCoq3, TgCoq5, TgCoq8, Arh1 homolog and TGGT1_232540 peptides. The data used to generate this volcano plot is included in **Supplementary Table 3**.

This approach allows us to identify proteins in close proximity to the baits and experimental validation would be required to demonstrate direct protein-protein interactions. We first validated that both *TgCoq3-TID* and *TgCoq5-TID* mutants were able to biotinylate mitochondrial proteins via IFAs (**Fig. 4B**). We performed a timepoint IFA by incubating the parasites with biotin for various lengths of time and saw labeling of the mitochondrion after 2 hours of biotin incubation (**Supplementary Fig. 6A**).

With the aim to find interacting proteins we performed immunoprecipitation using streptavidin magnetic beads with total lysates of the parental and *TgCoq3-TID* and *TgCoq5-TID* mutants previously incubated with biotin. The eluates were probed with αHA antibody, confirming successful pull down of both TgCoq3-TID and TgCoq5-TID (**Fig. 4C**, *top panel*). We also probed the elutes with αTgCoq3 which confirmed the presence of TgCoq3 in the TgCoq3-TID lysate as expected. Interestingly, TgCoq3 was also detected in the TgCoq5-TID pull down, validating the close proximity of these two proteins (**Fig. 4C**, *bottom panel*). Note that the size of TgCoq3 appears larger in the TgCoq3-TID eluate because of the addition of the TurboID tag, which increases the molecular weight by approximately 35 kDa. Mass spec analysis of the eluate of the *TgCoq5-TID* mutant (total lysate) revealed that only 20% of the enriched peptides (*TgCoq5-TID* elute/parental elute) were predicted to be mitochondrial by LOPIT^37^, while 25% were predicted to be cytosolic (**Supplementary Fig. 6B**). Since the UQ synthesis enzymes are predicted to be expressed at low levels, their signals may be masked by peptides labeled by immature TgCoq5-TID or by nonspecifically associated cytosolic proteins.

With the aim to increase enrichment of mitochondrial proteins and minimize cytoplasmic contamination, we developed a protocol that consisted in obtaining mitochondrial membranes enriched fractions of the TgCoq5-TID (+biotin), as it was expressed at higher levels than TgCoq3 (**Fig. 4C**). Additionally, we performed a similar subcellular fractionation of a cell line expressing TID in the cytosol. Because expression of TID by itself in the cytosol proved to be toxic to the parasites, we instead used a C-terminally tagged TID of the cytoplasmic protein FBXO14^43^.

Mitochondrial membrane enriched fractions (P2 as shown in the scheme shown in **Supplementary Fig. 4B**) were isolated from both cell lines and used as inputs for streptavidin IP. The elutes were analyzed by Mass Spectrometry, revealing high specificity of the approach, as 90% of the enriched peptides (fold change of peptides detected in TgCoq5-TID P2 over FBXO14-TID P2) were predicted to be mitochondrial (**Fig. 4D**). These enriched peptides are shown in red on the volcano plot (**Fig. 4E**). Importantly, the top hits from the pulldown (defined as those with more than a 4-fold enrichment and *p*-value <0.0001) were all predicted to be mitochondrial (**Table 1**). The most significantly enriched proteins included TgCoq3, TgCoq5, TgCoq8, and an Arh1-like protein, a putative UQ synthesis enzyme predicted to act as an electron carrier for Coq6 (**Supplementary Table 1**). We also identified several uncharacterized proteins predicted to be essential. The complete list of proteins obtained are shown in **Supplementary Table 3**.

**Table 1.**
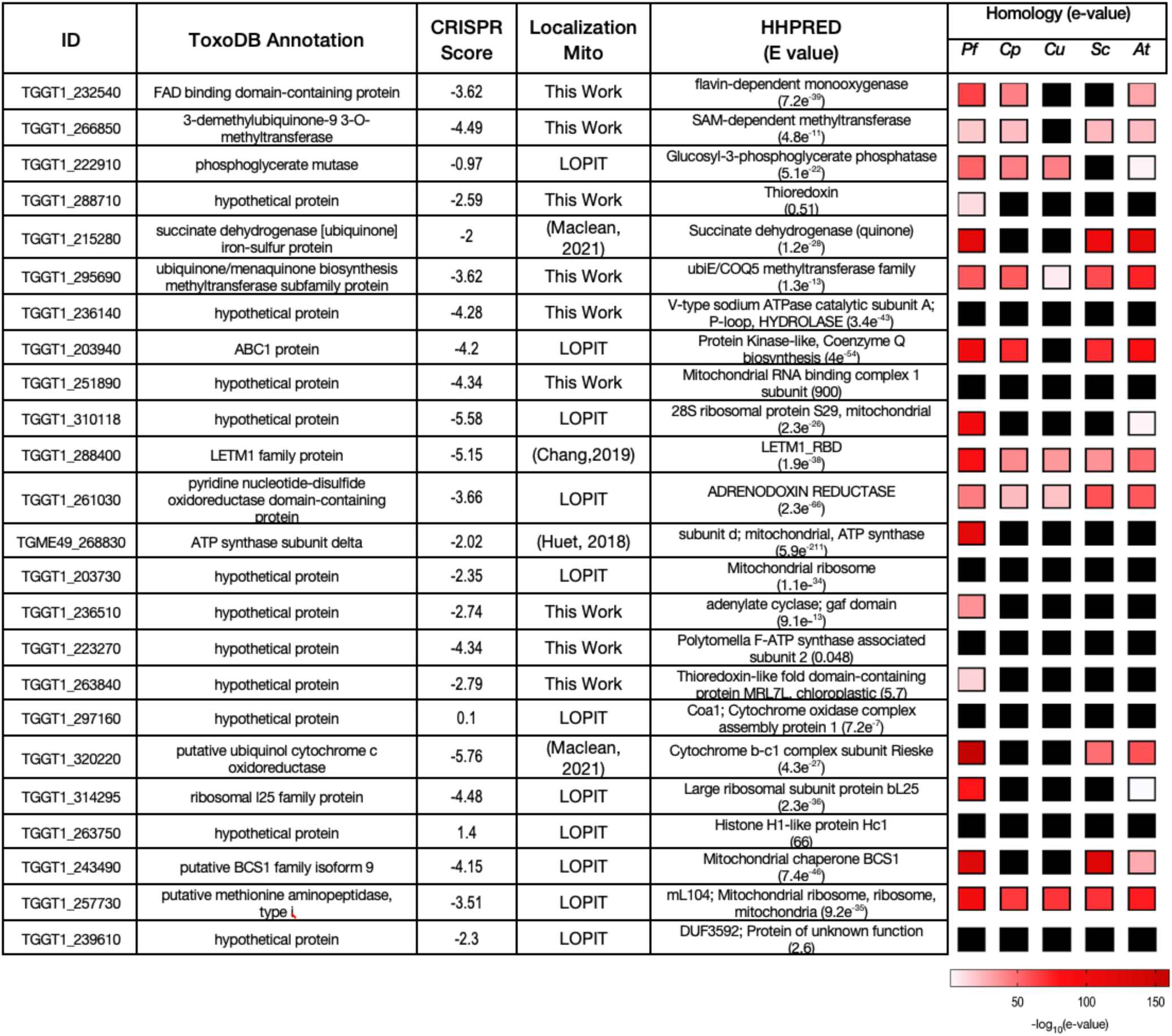
Proteins highly enriched in the *TgCoq5-TID* P2 fraction and their characteristics. List of proteins with *p* <0.0001 and fold change (TgCoq5-TID/ FbxO14-TID) higher than 4. GeneIDs, annotation in ToxoDB^36^, predicted localization from LOPIT^37^ (mm: mitochondrial membranes, soluble: mitochondrial lumen), and homologies of *Pf* (*Plasmodium falciparum*), *Cp* (*Cryptosporidium parvum*), *Cu* (*Cryptosporidium ubiquitum*), *Sc* (*Saccharomyces cerevisiae*), *At* (*Arabidopsis thaliana*) are shown. Black boxes indicate no homologous genes found.

In summary, we developed a protocol for enrichment of mitochondrial proteins for the identification of peptides proximal to TgCoq5. This strategy reduced nonspecific labeling by TurboID from ∼70% to less than 10%. This significant improvement enabled the discovery of several new candidates, likely part of the UQ synthesis pathway, which were undetectable in our initial experiments. We believe that combining biotinylation with subcellular fractionation offers a powerful strategy to uncover low abundance, functionally important proteins that are otherwise difficult to detect in total lysates.

### Discovery of a novel UQ modifying enzyme of plant origin that is absent in mammals

We reasoned that the combined proximity biotinylation and subcellular fractionation strategy would enable the identification of highly divergent proteins that form part of the UQ synthesis protein complex. To demonstrate this, we C-terminally tagged seven hypothetical proteins, identified by mass spectrometry and predicted to be mitochondrial, with a triple HA tag (**Supplementary** Fig. 7A), and confirmed their mitochondrial localization via IFA (**Supplementary** Fig. 7B). We next generated conditional knockdown mutants for the *TgCoq4* gene and four hypothetical genes that had a negative CRISPR score and were not predicted to be ribosomal proteins (**Supplementary Fig. 7C**). We first confirmed by plaque assays that two hypothetical genes, *TGGT1_288710* and *TGGT1_251890*, are essential for parasite growth, while *TGGT1_239610* and *TGGT1_236510*, predicted to be essential by CRISPR-based screens, were dispensable (**Supplementary Fig. 7D,E**).To assess their potential involvement in the UQ synthesis, we performed host cell lysis assays in a 96 well plates, culturing the mutants in media without ATc, with ATc, or with ATc and UQ_6_. The growth defect of the *iΔTgCoq4* mutant *(+ATc*) was rescued by UQ_6_ supplementation, validating the function of TgCoq4 in UQ synthesis. In contrast, none of the mutants deficient in the hypothetical genes showed rescued by UQ_6_ supplementation, indicating that although these proteins were biotinylated by TgCoq5-TID due to proximity, they are unlikely to be part of the UQ synthesis pathway (**Supplementary Fig. 7F**). One of the top hits from the mass spec analysis, TGGT1_232540 (hereafter referred to as TgCoqFAD), was annotated in ToxoDB^36^ as a “FAD binding domain containing protein”, and HHpred^35^ analysis of its sequence identified it as a “Flavin-dependent monooxygenase” (**Table 1**). We wondered if this gene and the corresponding protein could represent one of the missing activities for UQ synthesis, specifically Coq6, which is a flavin dependent monooxygenase in yeast and mammals. Based on bioinformatic analysis, TgCoqFAD possesses three FAD binding domains (**Fig. 5A**). In fungi and animals, the enzyme that carries out C6 hydroxylation is Coq7, a carboxylate-bridged diiron hydroxylase^44^. Interestingly, in plants, this enzymatic step is carried out by a flavin-dependent monooxygenase (FMO)^45^. By constructing a maximum likelihood tree, we discovered that TgCoqFAD is divergent from Coq6, Coq7, and bacterial UQ-hydroxylases, and is closely related to the plant FMO responsible for C6 hydroxylation. Moreover, other apicomplexans, such as *Plasmodium* (PF3D7_0815300), also possesses a homolog of TgCoqFAD, that is absent in yeast and mammals (**Fig. 5B**). Phylogenetic analysis using an expanded sampling of TgCoqFAD orthologs and paralogs revealed a clear pattern: alveolate sequences cluster with deltaproteobacteria (Desulfobacteria), while stramenopiles group with plants and green algae. In contrast, red algal sequences do not fall within this clade and instead group with more distant paralogs or outgroups (**Supplementary Fig. 8**).

**Figure 5.**
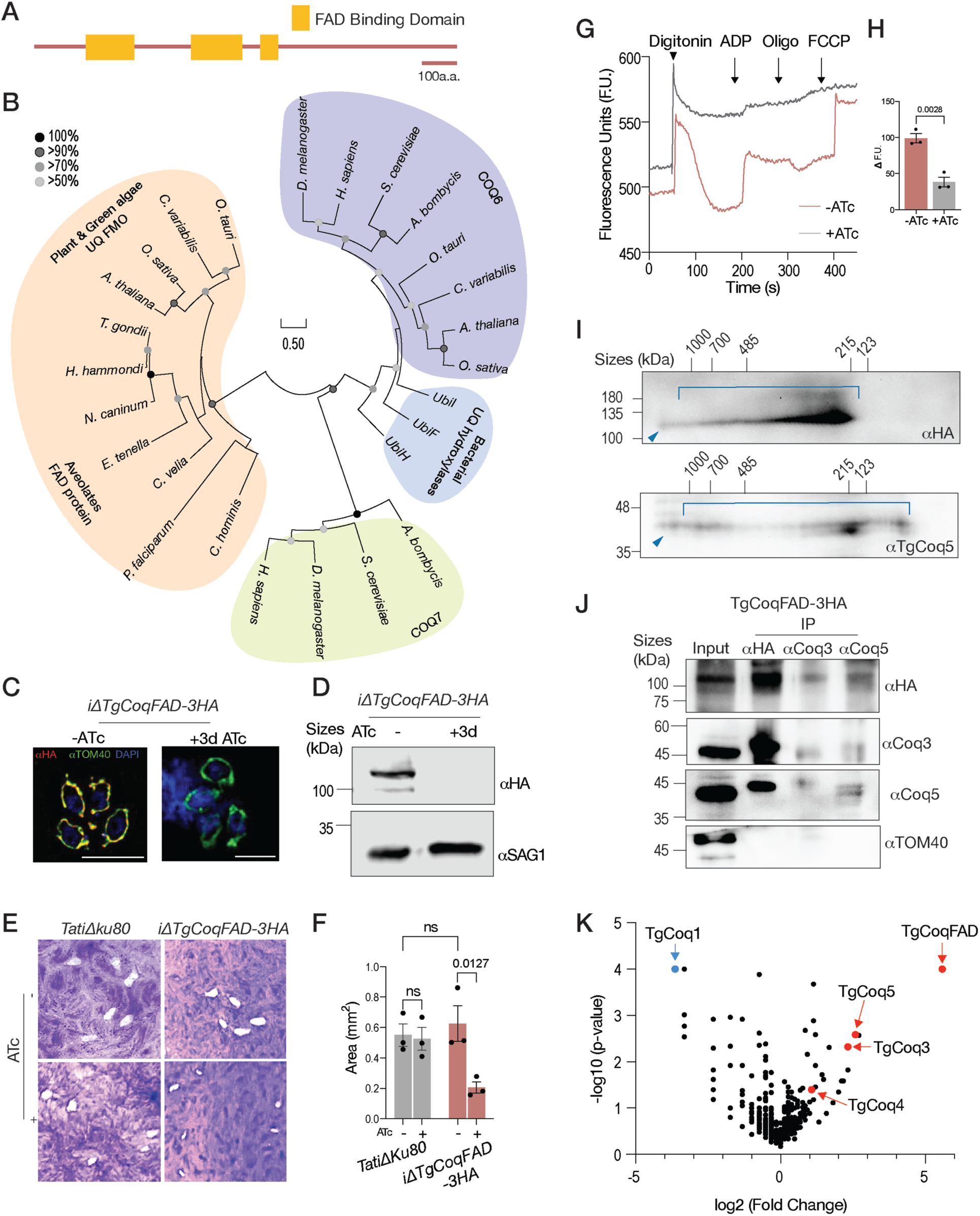
TgCoqFAD is essential for parasite growth and mitochondrial function and forms part of the UQ complex. (A) Predicted domains by Interpro of TGGT1_232540 (TgCoqFAD). (B) Phylogenetics of TgCoqFAD, UQ-hydroxylases, Coq7, and Coq6 from different organisms. Sequences are listed in **Supplementary Table 4**. (C) IFA showing TgCoqFAD localization to the mitochondrion using αTOM40 as mitochondrial marker and αHA to probe for TgCoqFAD. TgCoqFAD’s expression is downregulated by ATc. Scale bars represent 5 µm. (D) Western blot showing downregulation of TgCoqFAD with ATc incubation for three days. αHA was used to detect TgCoqFAD-3HA, and αSAG1 was used as loading control. (E) Representative plaques showing that knockdown of TgCoqFAD resulted in reduced plaque size. (F) Quantification of plaque areas from three biological replicates. Statistical significance was evaluated by two-way ANOVA. Each point represents a biological replicate, and error bars indicate SEM. (G) Representative tracing of membrane potential of *iΔTgCoqFAD* using Safranin *O*. <u>+ATc:</u> parasites were incubated with ATc for 3 days. <u>Oligo</u>: oligomycin. (H) *iΔTgCoqFAD* mitochondrial membrane potential quantification of the differences between maximum fluorescence (after depolarization with FCCP) and minimum fluorescence (after digitonin permeabilization). Each point represents a biological replicate, and the bars represent SEM. Student’s t test was used to evaluate significance, n=3. (I) 2D BN/SDS-PAGE using the *iΔTgCoqFAD-3HA* cell line lysate. The membrane was probed with αHA and αTgCoq5. Blue arrowhead indicates the UQ synthesis complex, and blue bracket indicates complex or disassembly intermediates. (J) Reciprocal co-IPs were performed using *iΔTgCoqFAD-3HA* lysates. TgCoqFAD-3HA, TgCoq3, and TgCoq5 were immunoprecipitated using αHA-, αTgCoq3-, and αTgCoq5-coupled beads, respectively. Equivalent amounts of input and IP eluates were analyzed by western blotting. αTOM40 was used as an IP control. See detailed protocol in the Methods section. The experiment was performed three times. (K) Volcano plot of peptides detected in the IP eluates plotted by fold change (TgCoqFAD-3HA/TgCoq1-3HA) and *p*-value. Red dots indicate peptides significantly enriched (at least 2-fold) in the eluate of *TgCoqFAD-3HA* that are involved in UQ synthesis. The blue arrowhead indicates TgCoq1, which was used as bait for TgCoq1-3HA IP. Data used to generate this volcano plot is included in **Supplementary Table 6**. For **F** and **H**, significance was considered when *p*-value is less than 0.05.

To characterize TgCoqFAD, we tagged its C-terminus with HA and generated conditional knockdown lines in the HA tagged background (*iΔTgCoqFAD-3HA*). We confirmed its mitochondrial localization (**Supplementary** Fig. 9A), and ATc regulation of expression via IFA and western blot analysis (**Fig. 5C and 5D**). To identify additional UQ synthesis enzymes of plant origin, we used the Arabidopsis Coq6 sequence (Q9LRM9) as a query and searched the *T. gondii* proteome using HHpred^35^. Interestingly, the top hits included TgCoqFAD and TGGT1_220360, the latter predicted to localize to the cytosol. HA tagging of TGGT1_220360 confirmed its cytosolic localization (**Supplementary Fig. 9A**).

To characterize the role of TgCoqFAD, we performed plaque assays in the presence and absence of ATc, revealing significantly smaller plaques with ATc treatment compared with the parental line (**Fig. 5E and 5F**). Lastly, we investigated the role of TgCoqFAD in mitochondrial activity. Downregulation of *TgCoqFAD* expression by ATc, caused a significant depolarization of the mitochondrial membrane in the mutant parasites (**Fig. 5G and 5H**). Importantly, using 2D BN/SDS-PAGE analysis of *iΔTgCoqFAD-3HA* whole cell lysates, we demonstrated that TgCoqFAD forms part of a large protein complex (**Fig. 5I**). We also probed the same membrane with αTgCoq5 and observed the same high MW complex (**Fig. 5I**). Note that the banding pattern differs from that observed in **Fig. 3B**, likely due to partial complex disassembly during sample preparation, as **Fig. 3B** was generated using the P2 mitochondrial fraction, while **Fig. 5I** was obtained from whole-cell lysates. Next, using reciprocal co-IPs, we validated the interactions between TgCoqFAD, TgCoq3 and TgCoq5 (**Fig. 5J**). By performing αHA IPs using the mitochondrial fractions from TgCoqFAD-3HA and TgCoq1-3HA (used as control), we saw TgCoq3, TgCoq4 and TgCoq5 being enriched when TgCoqFAD-3HA was used as bait (**Fig. 5K**), which further confirms the involvement of TgCoqFAD in the UQ synthesis complex.

### TgCoqFAD is a multifunctional monooxygenase essential for UQ synthesis

To validate the role of TgCoqFAD in UQ synthesis, we performed growth assays using *iΔTgCoqFAD-RFP* parasites. Culturing the mutant with ATc resulted in a significant growth defect (**Fig. 6A**, *+ATc*, gray line and bar). Importantly, supplementation of the media with UQ_6_ rescued this defect (*+ATc+UQ_6_*, purple line and bar), demonstrating that TgCoqFAD participate in the UQ synthesis pathway of *T. gondii* (**Fig. 6A**). Importantly, HPLC analysis of extracts of the *iΔTgCoqFAD* mutant (±ATc) demonstrated that UQ_7_ levels in the ATc treated parasites were reduced by more than ten-fold (**Fig 6B**), accompanied by the accumulation of the demethoxy-ubiquinone_7_ (DMQ_7_) precursor (**Supplementary Fig. 9B and 9C**). Together, these metabolic data provide direct biochemical evidence that TgCoqFAD is essential for UQ biosynthesis in *T. gondii*.

**Figure 6.**
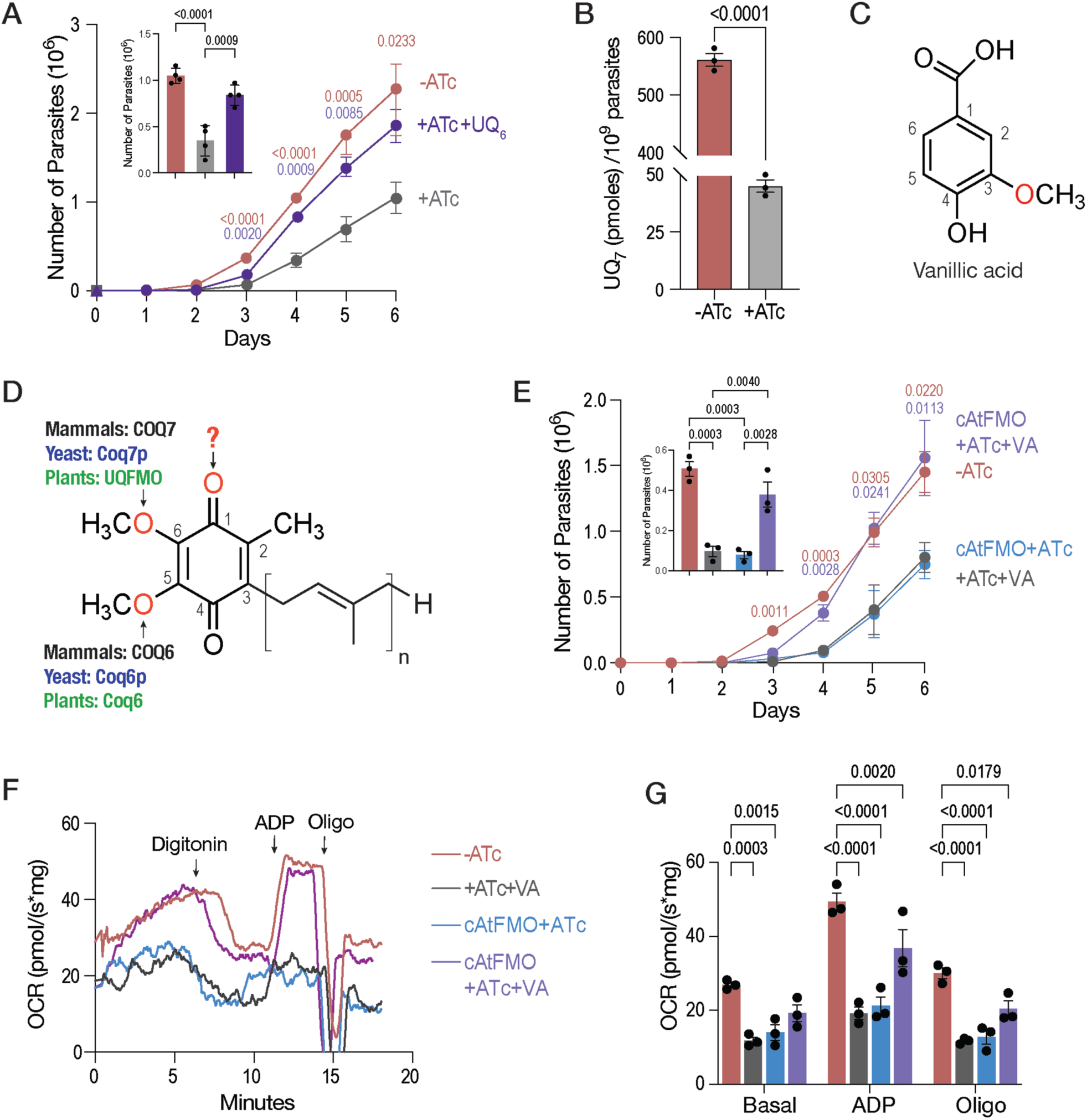
Genetic and chemical complementation reveal TgCoqFAD as a bifunctional monooxygenase. (A) Growth of *iΔTgCoqF*AD expressing RFP ± ATc or +ATc and 10 µM UQ_6_. Parasites were preincubated in ATc for three days prior to the infection. Data are presented as mean ± SEM. One-way ANOVA was used to evaluate significance, n=4. The bar graph shows the number of parasites at day 4. (B) Quantification of UQ_7_ from the *iΔTgCoqF*AD cell line incubated ± ATc (6 days). Data is presented as mean ± SEM. Student’s t test was used, n=3. (B) Structure of vanillic acid. (D) Structure of UQ_n_ and nomenclature of the eukaryotic hydroxylases involved in UQ ring decorations. (E) Growth of *iΔTgCoqF*AD-*RFP* complemented with the *Arabidopsis thaliana* FMO (cAtFMO). <u>-ATc</u>: *iΔTgCoqF*AD with ethanol control; <u>+ATc+10</u> <u>µM VA</u>: *iΔTgCoqF*AD grown with ATc supplemented with 10 µM Vanillic acid; <u>cAtFMO</u>: *iΔTgCoqF*AD complemented with AtFMO grown with ATc; <u>cAtFMO + 10 µM VA</u>: *iΔTgCoqF*AD complemented with AtFMO grown with ATc supplemented with 10 µM VA. Parasites were preincubated with ATc for three days prior to the infection. Data is presented as mean with SEM. Two-way ANOVA was used to evaluate significance, n=3. The bar graph represents the number of parasites at day 4. (F) Representative tracing of the oxygen consumption rate of the *iΔTgCoqFAD* mutants cultured using similar conditions to the ones used in **E** and presented using similar colors. (G) *iΔTgCoqFAD* mOCR quantification. Basal, ADP-stimulated, and oligomycin-inhibited mOCR were quantified from three biological replicates. Two-way ANOVA was used for statistical analysis, with the −ATc condition as the control group. Each point represents a biological replicate, and the bars represent SEM. For **A**, **B**, **E** and **G**, significance was considered when *p*-value is less than 0.05.

To investigate the enzymatic activity of TgCoqFAD, we used analogs of UQ precursors to bypass the effects of TgCoqFAD repression. Vanillic acid, which contains a methoxy group at the C3 position, required for C5 hydroxylation has been used to bypass COQ6 activity (**Fig. 6C**)^46^. Similarly, 2,4 dihydroxybenzoic acid, which is hydroxylated at both C2 and C4, required for C6 hydroxylation has been used to bypass COQ7 activity^47^. Although vanillic acid contains a methoxy group at C3 position, and 2,4-dihydroxybenzoic acid carries hydroxyl groups at the C2 and C4 positions, the C3 and C4 substitutions correspond to the C5 and C6 positions, respectively, in the final UQ ring after prenylation and decarboxylation. Consequently, supplying these compounds bypasses the requirement for COQ6 mediated C5 hydroxylation and COQ7 mediated C6 hydroxylation respectively (**Supplementary Fig. 9D**). However, neither precursor analog was able to rescue the growth defect caused by TgCoqFAD knockdown. This suggests that TgCoqFAD is not a monofunctional enzyme solely hydroxylating C5 or C6 position but rather may perform multiple enzymatic functions that cannot be bypassed by these modified ring precursors (**Supplementary Fig. 9E and 9F**). We also tested if 2,3,4-trihydroxybenzoic acid, which contains hydroxyl groups at C2, C3, and C4 positions, can bypass TgCoqFAD knockdown, however, this compound was toxic to the host cells and parasites were unable to grow (**Supplementary Fig. 9D-F**).

To confirm that the inability of benzoquinone analogs to bypass the pathway is not due to disassembly of the UQ complex upon TgCoqFAD knockdown, we assessed the expression levels of other UQ synthesis enzymes and evaluated the integrity of the UQ complex following TgCoqFAD depletion. Western blots and 2D BN/SDS-PAGE analysis, showed that expressions of TgCoq3 and TgCoq5 levels were not affected (**Supplementary Fig. 10A**). Moreover, the UQ complex is still present when knocking down TgCoqFAD (**Supplementary Fig. 10B**).

Since phylogenetic analysis indicates that *Arabidopsis thaliana* FMO (AtFMO) is the closest homolog to TgCoqFAD (**Fig. 5B**), and its function has been previously characterized^45^, we thought to complement the TgCoqFAD conditional knockdown cell line with AtFMO to infer the function of TgCoqFAD. This approach enables evaluation of their functional conservation in UQ synthesis. To ensure correct mitochondrial localization of AtFMO, we generated a plasmid encoding a chimeric protein consisting of TgCoqFAD’s mitochondrial targeting pre-sequence and a codon optimized mature sequence coding AtFMO. We validated the expression and mitochondrial localization of AtFMO (**Supplementary Fig. 10C**). However, growth kinetics revealed that AtFMO alone could not compensate for loss of TgCoqFAD, as the mutant parasites displayed the same growth defect in the presence of ATc (**Fig. 6E**, *cAtFMO+ATc*, *blue line and bar*). Interestingly, supplementing the growth medium of the *cAtFMO+ATc* mutant with vanillic acid fully restored its growth (**Fig. 6E**, *cAtFMO+ATc+VA*, *purple line and bar*). Vanillic acid provides a 3-methoxy group, which bypasses the requirement for the C5 hydroxylation step. Under these conditions, AtFMO, which catalyzes only the C6 hydroxylation^45^, becomes sufficient to support UQ synthesis. Full rescue of growth occurred only when AtFMO expression (providing C6 hydroxylation) was combined with vanillic acid supplementation (providing C5 modification) (**Fig. 6C and 6D**). These results demonstrate that TgCoqFAD normally performs both the C5 and C6 hydroxylation reactions during UQ biosynthesis. To further validate TgCoqFAD’s function, we measured mOCR in the mutants. Consistent with the mitochondrial membrane potential defect in the TgCoqFAD knockdown mutant, we observed a significant reduction of the mOCR when TgCoqFAD was depleted (**Fig. 6F and 6G**: *-ATc* and +*ATc*). Interestingly, in the *cAtFMO+ATc+VA* mutant, basal mOCR was fully restored, while ADP-stimulated and minimal mOCR (after oligomycin treatment) were increased relative to the +ATc condition but did not reach −ATc levels (**Fig. 6F and 6G**: *cAtFMO+ATc+VA*). Together, these functional data further support that TgCoqFAD catalyzes both C5 and C6 hydroxylation steps, identifying TgCoqFAD as the first bi-functional monooxygenase described in eukaryotic ubiquinone biosynthesis.

### TgCoqFAD can be targeted by small molecules in both tachyzoites and *in vitro* differentiated bradyzoites

The results presented highlight the discovery of TgCoqFAD, a unique enzyme absent in mammals that is integral to UQ synthesis, essential for mitochondrial function in *T. gondii*, and critical for parasite growth. These features underscore TgCoqFAD’s potential as a chemotherapeutic target. As a proof of concept for its targetability, we first predicted potential druggable pocket(s) on the AlphaFold-generated structure of TgCoqFAD using FTSite^48^. The latter identified few potential pockets that were proximal to the putative FAD binding site (**Suppleme**ntary Fig. 11). Using a GRID box large enough to encompass these pockets, we virtually screened a FDA-approved drug library (DrugBank approved library^49^) using FRED (OpenEye) which led to the identification of three structurally distinct compounds as best candidates for experimental validation given their predicted modes of interaction (**Fig. 7A and 7B**). Indeed, all three hits namely fenticonazole, flibanserin and redafamdastat inhibited parasite growth at low µM concentrations (**Fig. 7C**). To validate TgCoqFAD as the target of these compounds, we tested their EC_50_s in a TgCoqFAD overexpression line (TgCoqFAD-OE). Since the conditional knockdown line uses the SAG4 promoter, which is predicted to drive higher expression than the endogenous promoter^50^, we first confirmed the overexpression of TgCoqFAD in the *iΔTgCoqFAD* line via western blot (**Supplementary Fig. 12 A-B**). We observed more than a threefold increase in the EC_50_ values for fenticonazole and flibanserin, and over a fourfold increase for redafamdastat, in the TgCoqFAD-OE line, whereas the EC_50_ for atovaquone, used as control, remained unchanged. Moreover, host cells are still viable at 10X their respective EC_50_s, indicating the inhibition of parasite growth was not due to unhealthy host cells (**Fig. 7C**). Since the clinically used drug combinations, such as pyrimethamine and sulfadiazine, are only effective against tachyzoites, but lack activity against bradyzoites^51^, we next tested these compounds against *in vitro* differentiated bradyzoites. At 3X their respective EC_50_s, fenticonazole and redafamdastat significantly reduced bradyzoite viability (**Fig. 7D, 7E**, **and Supplementary Fig. 12 C**), supporting the targetability of TgCoqFAD in tachyzoites and its potential against bradyzoites. The limited activity of flibanserin against *in vitro*-derived bradyzoites may reflect reduced permeability into tissue cysts, an issue that will need to be addressed in future *in vivo* studies.

**Figure 7.**
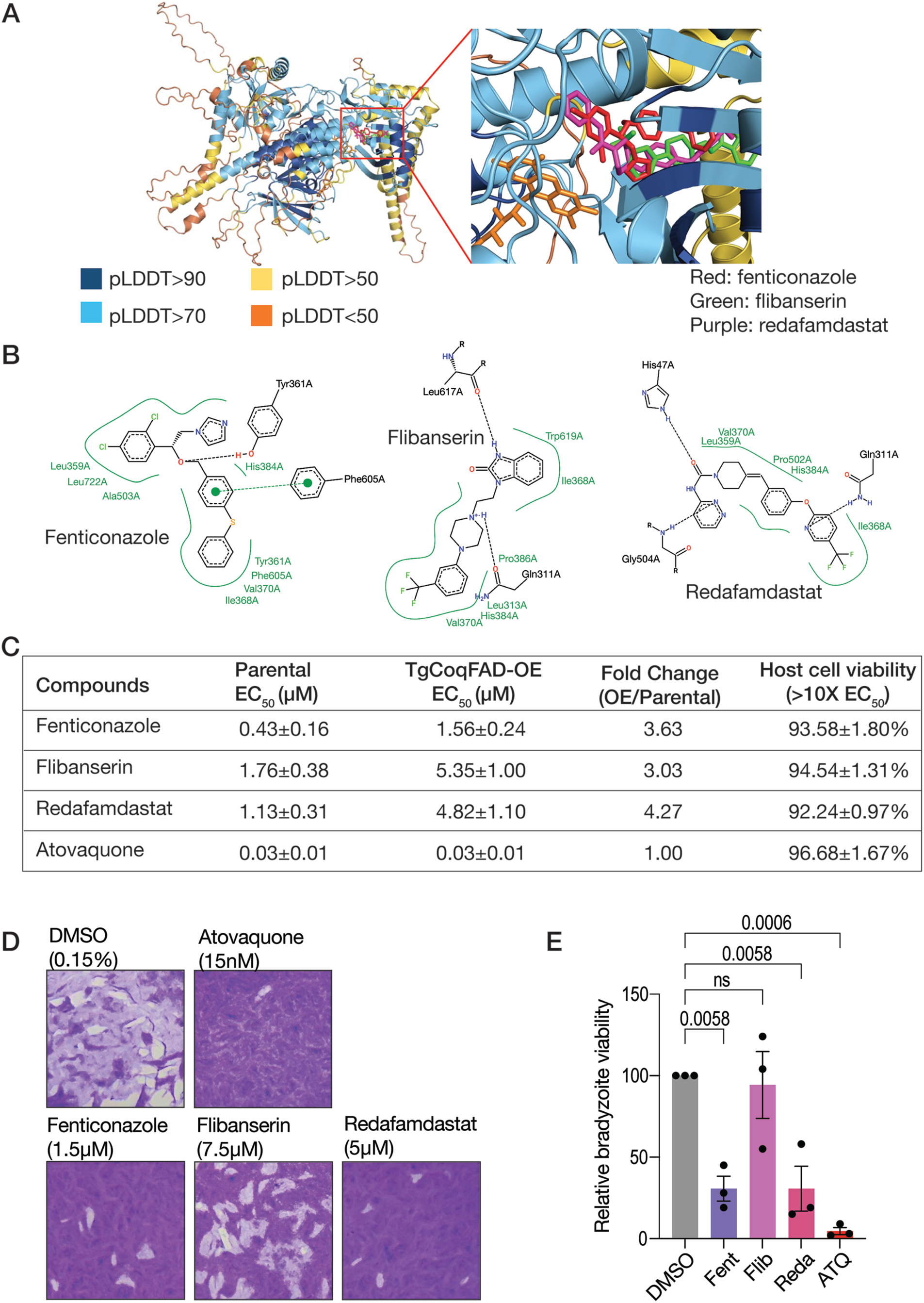
Molecular docking identifies TgCoqFAD inhibitors active against tachyzoites. (A) The TqCoqFAD model (uniprot id: A0A139Y5I3) generated shown in cartoon representation and colored based on the pLDDT score ranges shown. Global and close-up views of the predicted complex showing the proposed binding model of FAD and three DrugBank hits: fenticonazole, flibanserin, and redafamdastat, are shown on the left and right, respectively. All ligands are shown as sticks. FAD pose was modelled using AlphaFold3 while the poses of the three drugs are typical of the most reproducible and highest ranked poses predicted through GOLD-based focused docking done in 10 independent runs for each drug. The figure was generated in PyMol (open source, version 2.5). (B) 2D ligand interaction diagrams of the poses of the three drugs shown in A. These diagrams were generated using PoseView accessed through the Protein Plus server. (C) EC_50_ values of the compounds against parental (*TatiΔKu80* RFP), TgCoqFAD-OE (*iΔTgCoqFAD-3HA* RFP), and fold change between TgCoqFAD-OE over parental. Cytotoxicity was tested at 10 X EC_50_ obtained from the parental line. Assay was conducted with 3 biological replicates each containing at least 2 technical replicates or 3 technical replicates for cytotoxicity. EC_50_s were presented as mean ± SEM. (D) Representative images showing the viabilities of the bradyzoites treated with 3X EC_50_ of the inhibitors. DMSO and Atovaquone treatments were used as negative and positive controls respectively. (E) Bar graph shows the quantification of relative bradyzoite viability, percentage of plaques normalized to DMSO treatment. Fent: fenticonazole, Flib: flibanserin, Reda: redafamdastat. Data represented as mean with SEM, n=3. One-way ANOVA test used. Significance was considered when *p*-value was less than 0.05.

In summary, these data support the potential of TgCoqFAD as a therapeutic target. Although the three compounds we tested exhibited EC_50_ values that are not outstanding, they nonetheless provide proof of concept that TgCoqFAD, an enzyme absent in mammals, could be effectively targeted for therapeutic intervention.

## DISCUSSION

In this work, we characterized key components of the ubiquinone (UQ) synthesis pathway in *Toxoplasma gondii*, identifying TgCoq3, TgCoq4, TgCoq5, TgCoq8, and a novel bifunctional flavin-dependent monooxygenase, TgCoqFAD, which is present in apicomplexan parasites but absent in mammals and yeast. We show that these proteins assemble into a mitochondrial UQ synthesis complex. Conditional knockdown of each component impaired parasite growth and mitochondrial activity, and the growth defects were rescued by exogenous UQ_6_ supplementation, validating their roles in UQ biosynthesis. Interestingly, *iΔTgCoq3, iΔTgCoq5*, and *iΔTgCoqFAD* were still able to proliferate following conditional knockdown, though at a significantly slower rate. This is most likely due to the long half-life of UQ^52,53^ and its ability to be recycled through the Q cycle, allowing residual UQ to support minimal growth even after knockdown. A key function of UQ is to shuttle electrons from various dehydrogenases (such as complex II) to complex III, which then transfers electrons to cytochrome c in the mitochondrial ETC. In agreement with this role, *iΔTgCoq3*, *iΔTgCoq5* and *iΔTgCoqFAD* showed mitochondrial dysfunction, as evidenced by depolarization of their mitochondrial membrane potential and reduced oxygen consumption rates. Unlike other organisms, the *T. gondii* UQ synthesis complex migrate at over 1 MDa by two-dimensional BN/SDS-PAGE analysis, which is larger than the yeast UQ synthome (∼690 kDa) and the mammalian complex Q (less than 480 kDa). This size difference resembles the distinctive features of other *T. gondii* mitochondrial complexes, which are larger compared to the yeast and human counterparts^32,54^.

TgCoq5 was previously detected in the complexome profiling study by Maclean et al.^32^, however, none of the other UQ synthesis proteins were detected, likely due to their low abundance. The use of the biotin ligase^42^ approach facilitates the identification of low-abundance proteins but it is prone to false positives, especially for mitochondrial proteins. Most mitochondrial proteins are encoded in the nucleus^55^, and they transiently associate with cytosolic proteins prior to their import into the mitochondrion. To overcome this limitation, we combined biotinylation with subcellular fractionation to separate mitochondrial proteins from cytosolic contaminants and included a cytosolic TurboID as a spatial control. This strategy significantly improved the mitochondrial yield from 20% in total lysates to 90% in mitochondrial fractions.

Our mass spec results revealed that TgCoq5-TID is in proximity to TgCoq3, Coq8 and Arh1 homologs. Although TgCoq4 was not detected in the proximity labeling dataset, we independently validated its role in UQ synthesis: conditional knockdown of TgCoq4 caused a growth defect that was rescued by UQ_6_ supplementation, and 2D BN/SDS-PAGE analysis confirmed its association with the UQ complex. We hypothesize that TgCoq4 function as an oxidative decarboxylase similar to its mammalian homolog COQ4^56^. Since the UQ complex exceeds 1,000 kDa, we hypothesize that the biotin ligase fused to TgCoq5 could be oriented away from TgCoq4 limiting its biotinylation. Another explanation could be that the peptide signal is masked by more abundant subunits of the ETC complex.

Interestingly, several UQ synthesis components that are part of the canonical CoQ synthome in yeast, including Coq6, Coq7, Coq9, and Coq11, do not have identifiable homologs in *T. gondii*. In yeast, Coq9 facilitates Coq7 activity, and Coq6 and Coq7 together carry out essential hydroxylation steps required for Coq3-mediated *O*-methylation. In contrast, our data indicate that *T. gondii* has replaced these activities with the bifunctional enzyme, TgCoqFAD, which performs the roles of both Coq6 and Coq7. In this context, the absence of Coq9 in *T. gondii* likely reflects functional streamlining of the pathway.

Although the *T. gondii* UQ synthesis complex is larger in apparent molecular weight than those described in other organisms, it is composed of fewer distinct protein components, highlighting an evolutionary reorganization of the UQ synthesis machinery in apicomplexan parasites. TgCoqFAD lacks clear homologs in yeast or mammals, explaining why it was not detected in previous bioinformatic analyses. Phylogenetic analysis revealed that TgCoqFAD is most closely related to plant flavin-dependent monooxygenase (FMO) known to catalyze C6 hydroxylation. Previous work on plant FMOs suggests that the evolutionary branching occurred via horizontal gene transfer from a bacterium harboring the ancestral gene to green algae^45^. Considering that apicomplexans and plants both belong to Diaphoretickes, whereas animals and fungi belong to Amorphea^57^, it is not surprising that *T. gondii* utilizes an enzyme homologous to plant FMO for this essential mitochondrial pathway. However, this evolutionary pattern is not consistent with inheritance from the red algal endosymbiont that gave rise to the apicoplast and instead suggests that TgCoqFAD was likely acquired through other evolutionary routes, such as horizontal gene transfer. This raises the intriguing possibility that additional metabolic enzymes in *T. gondii* may have been acquired in a similar manner, contributing to the unique organization of its mitochondrial pathways.

Using comparative analysis of mitochondrial proteomes, we identified only 25 mitochondrial proteins of presumed plant origin, including TgCoqFAD. Replacement of the iron-dependent Coq7 with an FAD-dependent monooxygenase may provide a selective advantage by reducing dependence on labile iron and limiting iron-catalyzed reactive oxygen species (ROS) production^58,59^. In *T. gondii*, which faces fluctuating iron availability during its life cycle, such an adaptation may help maintain mitochondrial function. In *Plasmodium spp*., despite replication within heme-rich red blood cells, iron is tightly regulated, representing a major source of oxidative stress rather than a freely available cofactor^60^. Thus, replacing the iron-dependent Coq7 with FMO could provide significant advantages for apicomplexan parasites. Because TgCoqFAD is absent in the host and essential for parasite replication, it represents an attractive chemotherapeutic target for apicomplexan diseases such as toxoplasmosis and malaria. Moreover, although ATP synthase is dispensable in blood stage *Plasmodium* parasites^61^, UQ remains essential throughout all stages of the parasite’s life cycle. Apart from its role in oxidative phosphorylation, UQ also functions as an electron acceptor for dihydroorotate dehydrogenase, a key enzyme in the essential pyrimidine biosynthesis pathway^62^. UQ’s involvement in multiple critical biological processes, combined with the parasite inability to use host UQ_10_^30^, further supports TgCoqFAD as a promising drug target.

To investigate whether a second FMO might participate in UQ synthesis, we queried the *T. gondii* proteome using the plant Coq6 sequence (Q9LRM9). The top hits were TgCoqFAD and TGGT1_220360, which did not localize to the mitochondrion. While all eukaryotic FMOs involved in UQ synthesis, including the plant FMO, are strictly mono-functional, bi- and tri- functional monooxygenases, such as UbiL and UbiM, are present in certain bacteria species, providing precedent for a single enzyme catalyzing multiple hydroxylation steps (**Supplementary** Fig. 1). Interestingly, UbiL and UbiM are found in genomes that encode multiple UQ-hydroxylases, while in genomes encoding a single UQ hydroxylase, UbiM is typically the sole enzyme present, supporting a model in which bifunctionality compensates for reduced pathway complexity. UbiM has been identified in alpha-, beta-, and gamma-proteobacteria, but it is less distributed and conserved than other UQ-hydroxylases, and its origin remains a mystery^58^. The bioinformatic analysis, combined with the results of the genetic and chemical complementation experiments, provides clear evidence that TgCoqFAD catalyzes both the C5 and C6 hydroxylation steps of UQ biosynthesis. The inability of plant AtFMO to rescue growth unless vanillic acid is supplied demonstrates that TgCoqFAD performs a function that cannot be replaced by a mono-functional FMO. We propose that TgCoqFAD originated from an ancestral UbiM-like enzyme acquired through horizontal gene transfer.

Our results also underscore the therapeutic potential of TgCoqFAD. Virtual screening guided by an AlphaFold predicted model identified structurally diverse compounds with strong predicted binding affinity for TgCoqFAD. The *in vitro* efficacy of these compounds, combined with their reduced potency in a TgCoqFAD overexpression line, provides functional evidence that they likely act on this target. Importantly, the specificity of this effect, absent in a control treatment with atovaquone^12^, reinforces the idea that TgCoqFAD is not only essential for parasite long-term growth but also selectively druggable. This positions TgCoqFAD as a compelling target for future drug development efforts aimed at treating toxoplasmosis and potentially other apicomplexan infections.

Despite these advances, we were unable to directly assess TgCoqFAD’s enzymatic activity *in vitro*. This limitation is shared with many UQ biosynthesis enzymes, which often require integration into multiprotein complexes for catalytic function^63^. Reconstituting such complexes under *in vitro* conditions remains challenging. Additionally, while AlphaFold based molecular docking offers a valuable starting point for structural insights, the inherent uncertainty of predicted models must be acknowledged. Therefore, these docking results should be interpreted as preliminary and serve as proof of concept of TgCoqFAD’s potential clinical relevance. Future work aimed at resolving high resolution cryoEM or crystal structure of TgCoqFAD will be critical for advancing drug design. Furthermore, there are currently no studies investigating gene essentiality in the bradyzoite stage. Our *in vitro* bradyzoite drug testing data suggest that TgCoqFAD may represent a promising target in this stage of the parasite. However, additional work will be required to confirm the essentiality of the UQ synthesis genes for bradyzoites.

Interestingly, some azoles have been reported to have modest anti-*Toxoplasma* activity^64^, although no studies have previously shown that fenticonazole inhibits apicomplexan parasites growth, making this study the first indication of its activity. It is important to note that these compounds might have pleiotropic effects. Future medicinal chemistry efforts and target-focused compound optimization could build on these initial hits to develop more potent and specific TgCoqFAD inhibitors.

In summary, we uncover the unique UQ biosynthesis machinery in *T. gondii*, a key component of the parasite mitochondrial function (**Fig. 8**). We identified key components of the UQ complex, TgCoq3, TgCoq4, and TgCoq5, and characterized TgCoqFAD, an apicomplexan-specific bi-functional flavin-dependent monooxygenase catalyzing both C5 and C6 hydroxylation. To our knowledge, this is the first example of a bi-functional UQ monooxygenase in a eukaryotic system. This unprecedented bifunctionality likely provides an evolutionary advantage in coping with iron fluctuations and redox stress. Importantly, TgCoqFAD is essential for parasite long-term proliferation, absent in hosts, and druggable, positioning it as a highly promising antiparasitic target. Our findings lay the ground for developing targeted therapies against toxoplasmosis, malaria, and related diseases.

**Figure 8.**
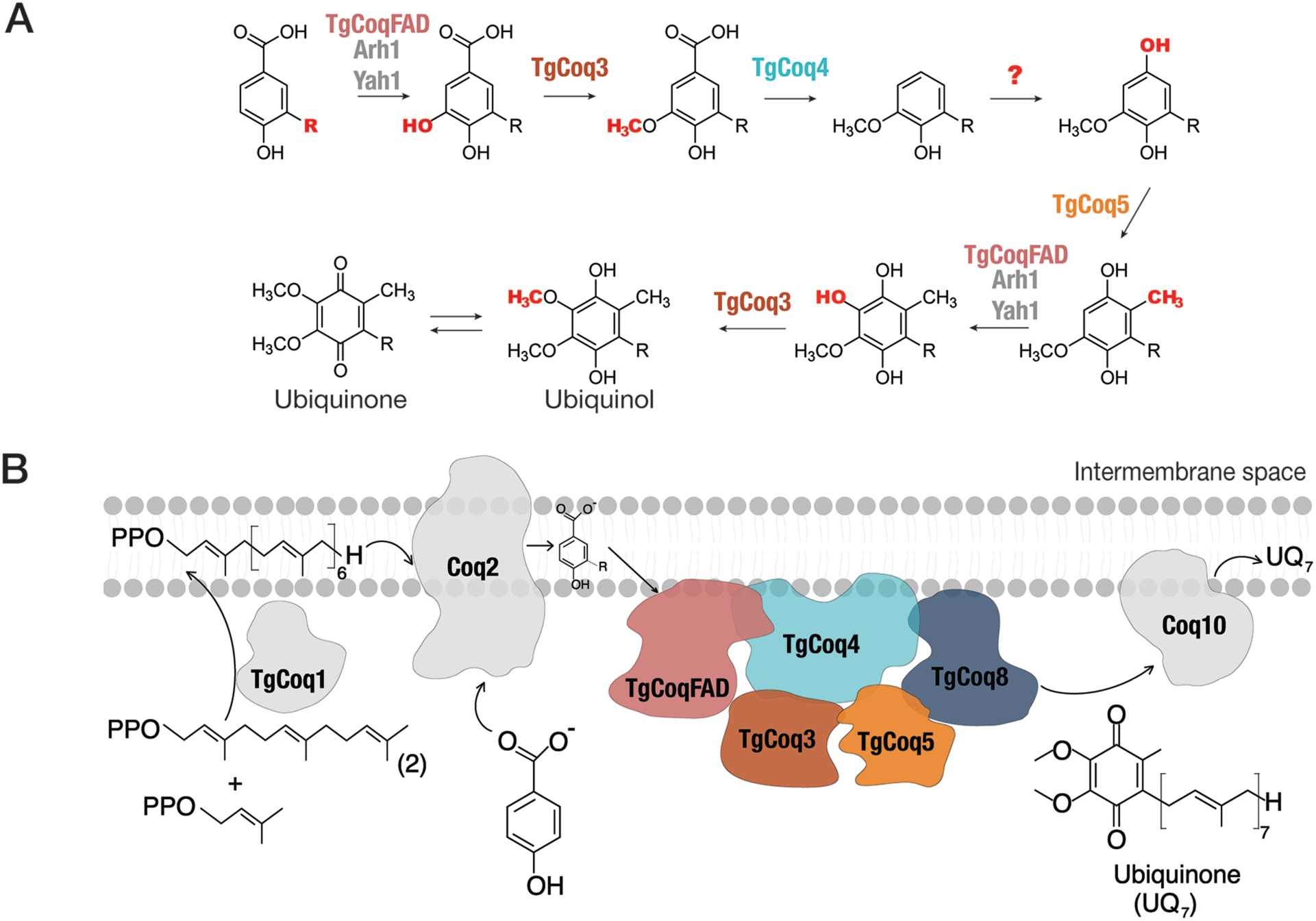
Proposed model for UQ biosynthesis and complex organization in *T. gondii.* (A) Proposed aromatic ring modification steps of UQ, highlighting the enzymes characterized in this study (TgCoq3, TgCoq4, TgCoq5, and TgCoqFAD) and the proposed electron donors for TgCoqFAD (Arh1 and Yah1 homologs). Question marks indicate steps that have not been experimentally determined. (B) Proposed model of the *T. gondii* ubiquinone synthesis complex.

## Acknowledgement

We thank the CTEGD FACS sorting and biomedical microscopy cores for the use of their facilities and equipment. Julie Nelson and Muthugapatti K Kandasamy provided training and support. We thank the Proteomics & Metabolomics Facility (RRID:SCR_021314), at the Nebraska Center for Biotechnology at the University of Nebraska-Lincoln for the mass spectrometry analysis. The facility and instrumentation are supported by the Nebraska Research Initiative. We thank John Smith, a veterinary scholar for performing the initial BLAST search during the summer of 2020. We thank Melissa Ann Sleda for initial training on the use of the established *Toxoplasma* protocols, Mayara Bertolini for training on the use of the Oroboro and protein purification, Jessica Kissinger for help with bioinformatics, Elisabet Gas Pascual from Chris West lab for training on co-immunoprecipitation, Madelaine Usey for training on BlueNative PAGE. Andrea Hortua Triana directed the generation of the αTgCoq3 and αTgCoq5 antibodies. The *FbxO14-TID* cell line was a gift from the Diego Huet lab. The αTOM40 antibody was a gift from Giel van Dooren, and the monoclonal αHA antibody was a gift from Chris West. We thank Justin Wiedeman from Etheridge lab for sharing his protocol for the elution of biotinylated proteins from streptavidin beads.

TR acknowledges OpenEye (Cadence Design Systems) for granting academic license for Omega and FRED (OEDocking).

This work was funded by NIH grant AI169846 to S.N.J.M and NSF grant MCB-2216747 to G.J.B. B.B. was partially funded by AHA predoctoral fellowship 24PRE1192541.

## Author contributions

B.B performed most of the experiments, analyzed the data and wrote the manuscript. T.R. performed molecular docking, virtual screening, and edited the manuscript. S.L. and G.J.B performed lipid extraction and quinone measurements via HPLC and edited the manuscript. S.N.J.M. supervised the project and wrote and edited the manuscript.

## METHODS

### Culture

hTERT (human telomerase reverse transcriptase) cells were cultured in Dulbecco’s modified minimal essential medium (DMEM) supplemented with 10% BCS (Gemini Bio-Products, 100-506-500). Cell cultures were maintained in 5% CO_2_ at 37°C. *T. gondii* tachyzoites were cultured in hTERT cells in DMEM with 1% BCS and purified as described previously^30^.

The C-terminally tagged line mixed populations were cultured in DMEM with 1% BCS with 6.8 µg/mL Chloramphenicol (Sigma, C0378). The conditional knockdown mixed populations were cultured in DMEM with 1% BCS containing 1 µM pyrimethamine (Sigma, P7771). Complementation lines mixed populations were cultured in DMEM with 1% BCS with 10 µM fluorodeoxyuridine (Sigma, F0503). The clonal lines were cultured in DMEM with 1% BCS. For downregulation of the genes, 0.5 µg/mL anhydrotetracycline (ATc) (Cayman Chemical Co Inc, 10009542-500) was added.

### Phylogenetic analysis

Protein sequences were obtained from the National Center for Biotechnology Information (NCBI)^34^ database. The *T. gondii* TgCoq3 (TGGT1_266850), TgCoq5 (TGGT1_295690) and TgCoqFAD (TGGT1_232540) sequences were used to query the NCBI database using the genomic Basic Local Alignment Search Protein (BLASTP) tool by selecting organisms in the apicomplexan phylum. Other sequences for Coq3 and Coq5 were obtained through UniProt^65^ by searching Coq3 and Coq5 in eukaryotes and UbiG and UbiE in prokaryotes. The phylogenetic tree from Latimer et al.^45^ was used as a guide to search for Flavin-dependent monooxygenases in plants and green algae. Coq7 sequences were obtained by searching the sequences in UniProt. For supplementary Fig. 8, sequences were obtained from BLASTP using Blossum45 setting.

The evolutionary history was inferred by using the Maximum Likelihood method and JTT matrix-based model^66^. The bootstrap consensus tree inferred from 100 replicates^67^ is taken to represent the evolutionary history of the taxa analyzed^67^. Branches corresponding to partitions reproduced in less than 50% bootstrap replicates are collapsed. The percentage of replicate trees in which the associated taxa clustered together in the bootstrap test (100 replicates) are shown next to the branches^67^. Initial tree (s) for the heuristic search were obtained automatically by applying Neighbor-Join and BioNJ algorithms to a matrix of pairwise distances estimated using the JTT model and then selecting the topology with superior log likelihood value. Evolutionary analyses were conducted in MEGA X^68^.

### Generation of mutant parasites

For C-terminus tagging, we used CRISPR/Cas9-based genome editing^69^ to introduce a 3xHA tag and a chloramphenicol drug resistant cassette into the 3’ end of the gene of interest (GOI) upstream to the stop codon. We used a single guide RNA targeting the 3’ end of the open reading frame into pSAG1-CAS9-U6-sgUPRT^70^ using Q5-site directed mutagenesis (NEB, M0491L). The primers used for all mutants are listed in the **Supplementary Table 7,** and we confirmed the insertion by Sanger sequencing. For each GOI, we amplified a donor DNA sequence encoding 3xHA from pLic-3HA-CAT^39,71^ in tandem with 40 bp flanks of the GOI as listed in **Supplementary Table 7**. For generation of TurboID tagged lines, we amplified a donor DNA sequence encoding TurboID-3HA from the 3xHA TurboID pLIC plasmid (modified from Addgene #107169) in tandem with 40 bp flanks of the 3’ end of TgCoq3 or TgCoq5.

For generation of conditional knockdown cell lines, we used a single guide RNA targeting the 5’ open reading frame of the GOI into either pSAG1-CAS9-U6-sgUPRT^72^ using Q5-site directed mutagenesis or the pSS013^69^ using restriction enzymes as previously described^69^. The primers used are listed in **Supplementary Table 7**, and the sequences were confirmed by sequencing. For each GOI, we amplified the donor DNA sequence encoding Tet7SAG4 and a pyrimethamine drug resistant cassette from the pDTS4myc plasmid^39^ in tandem with 40 bp flanks of the GOI using primer pairs listed in **Supplementary Table 7**. For each cell line, we transfected the modified CAS9 plasmid and donor DNA by electroporation into *TatiΔku80* or the C-terminus tagged line as mentioned in the results.

For generation of the complementation cell lines, we used a single guide RNA targeting the UPRT locus using pSAG1-CAS9-U6-sgUPRT^72^ and amplified donor DNA sequence encoding SAG4 promoter, TgCoqFAD full length cDNA without stop codon (synthesized by Twist Bioscience, TGGT1_232540), 3xTy tag and a stop codon in tandem with 40 bp flanks of UPRT. For complementation with AtFMO, we cloned codon optimized AtFMO (synthesized by Twist Bioscience, UniprotID: Q8GYJ7) into pSAG4-TgCoqFAD-3Ty plasmid using primers listed in **Supplementary Table 7**. To ensure correct localization, we retained the endogenous mitochondrial targeting sequence (MTS) from TgCoqFAD and removed the plant MTS from the AtFMO sequence. Transfected parasites were selected with 10 µM fluorodeoxyuridine.

The generation of RFP expressing parasites was done as described^73^. In brief, the plasmid expressing tdTomato^74^ was transfected into the conditional knockdown mutants. The mixed populations were sorted two times using a Bio-RadS3 cell sorter selecting for red fluorescent parasites. Positive clones were obtained through serial dilution.

### Cloning and antibody generation

cDNA from *TgCoq3* (TGGT1_266850) and *TgCoq5* (TGGT1_295690) was cloned into the pQE80L (Qiagen) plasmid and transformed into *E. coli* BL21 codon plus. Protein expressions were induced with 1 mM IPTG, and both recombinant proteins were purified from the insoluble fraction. For TgCoq3, 100 µg of purified protein was first mixed with Freund’s complete adjuvant and the mix used to inoculate mice (intraperitoneal) followed by three boosts 2 weeks apart with 50 µg of protein in Freund’s incomplete adjuvant^75,76^. For TgCoq5, 200 µg of purified protein was first mixed with Freund’s complete adjuvant and used to inoculate guinea pigs subcutaneously followed by 3 boosts 3 weeks apart, each with 100 µg of protein in Freund’s incomplete adjuvant. The final serums from mice and guinea pigs were tested by western blots with total parasite lysates and validated with lysates of the corresponding conditional knockdown cell lines (validation is shown in supplementary Figs. 3C and 3F). Both antibody serums were affinity purified prior to use for IFAs and westerns. Work with mice and guinea pigs were carried out in strict accordance with the Public Health Service Policy on Humane Care and Use of Laboratory Animals and Association for the Assessment and Accreditation of Laboratory Animal Care guidelines. The animal protocol was approved by the University of Georgia’s Committee on the Use and Care of Animals (protocol A2021 03-005-A5). All efforts were made to humanely euthanize the mice and guinea pigs after collecting blood.

### Immunofluorescence assay (IFA) and western blots

For IFAs with intracellular tachyzoites, sub-confluent hTERT cells seeded onto coverslips were infected for 20-24 h and the coverslips fixed with 3% paraformaldehyde for 15 min and permeabilized with 0.25% Triton X100 for 10 min. The slides were blocked in PBS pH 8.0 containing 3% bovine serum albumin (BSA) for 30 min prior to incubation with the respective antibody. Primary and secondary antibodies were incubated for 1 h each. Antibodies were diluted in PBS pH 8.0 with 3% BSA. Primary purified mouse αTgCoq3 was used at 1:25, purified guinea pig αTgCoq5 was used at 1:500, rabbit αTOM40^38^ (gift from the van Dooren lab) was used at 1:20,000. Secondary antibodies include GaM CW800 (LI-COR, 925-32210), GaGP 680RD (Neta Scientific, LIC-925-68077), GaRab HRP (Bio-Rad, 1706515), GaM HRP (Invitrogen, 31430), GaGP HRP (Invitrogen, A18769) and Streptavidin CW800 (LI-COR, 926-32230), were used at 1:1,000. Coverslips were mounted on glass slides with DAPI fluoromount-G. Images were taken with a Delta Vision microscope (Olympus, IX-71) with a Photometric Coolsnap camera. Deconvolved images were obtained by using softWoRx deconvolution software.

For western blots, PAGE separated proteins were transferred onto nitrocellulose membranes and blocked in 5% non-fat milk in PBS-T (0.1% Tween-20 in PBS) for 30 min. Incubation with primary antibodies was done overnight at 4°C, and secondary antibodies for 1 hour at room temperature. Antibodies were diluted in PBS-T. Primary purified mouse αTgCoq3 was used at 1:250, primary purified guinea pig αTgCoq5 was used at 1:500, αHA (bioLegend) was used at 1:1,000, αHA (gift from Christopher West’s lab) at 1:10,000, mouse αTubulin at 1:30,000 (Sigma, T6199), mouse αF1β (Agrisera) at 1:10,000, and mouse αSAG1^77^ (gift from the Carruthers lab) was used at 1: 10,000. The protein bands were imaged with a Licor Odyssey CLx equipment (LI-COR, 9140) using secondary antibodies that are fluorescent at either 680 or 800 nm, or imaged with ChemiDoc system (Bio-Rad, ChemiDoc^TM^ Imaging system) using HRP secondary antibodies.

### Lytic cycle assays, UQ_6_ rescue, and analog bypass

hTert cells were seeded onto 6-well plates and cultured until confluent. 150 parasites were plated onto the wells for each condition. The parasites were incubated for 7 days without disruption in the media with or without ATc. At day 7, infected host cells were fixed with 100% ethanol and stained with gentian violet as previously described^78^. ImageJ software was used to measure the size of the plaques. 30 plaques were randomly selected and measured for each condition and averaged for each biological repeat.

For the 96 well cell lysis assay, hTert cells were seeded onto 96 well plates and cultured until confluent. 4,000 parasites were plated in each well in triplicate. The parasites were incubated for 5 days in the media either without ATc, with ATc, or with 10 µM UQ_6_ and ATc. Infected host cells were then fixed and stained as described above and the absorbance of gentian violet was measured at 592 nm.

Replication assays were carried out as previously described^79^. hTert cells were seeded on coverslips in 12-well plates and cultured until sub-confluent. 5 x 10^5^ RFP-expressing parasites were used to infect the host cells and cultured in media either with or without ATc for 24 h before fixing using the same protocol as for IFA. The Delta Vision microscope was used for counting parasites per vacuoles. Two coverslips were infected per condition, and 100 parasitophorous vacuoles were randomly selected per coverslip. A total of three biological experiments were conducted.

Growth assay experiments were carried out using parasites expressing tdTomato (red fluorescent protein)^80^ with a previously described modification^81^. RFP-expressing parasites were preincubated with ATc for two days prior to the growth assay. Preincubated and non-preincubated parasites were used to infect confluent hTert cells in 96-well plates. Parasites were cultured in clear DMEM media either without ATc, with ATc, with ATc plus 10 µM UQ_6_, or with ATc plus vanillic acid or 2,4 di-hydroxybenzoic acid (concentrations as indicated in the figure legend). Fluorescence from RFP was read daily with a Biotek synergy H1 plate reader (Agilent, H1MFDG). Standard curves were used to convert the fluorescence values to number of parasites^73^.

### Mitochondrial membrane potential and oxygen consumption

*iΔTgCoq3*, *iΔTgCoq5*, and *iΔTgCoqFAD-3HA* were used to infect confluent hTert in T75 flasks and cultured in media with or without ATc for two days. Freshly egressed parasites were collected, washed with BAG (116 mM NaCl, 5.4 mM KCl, 0.8 mM MgSO_4_, 50 mM HEPES pH 7.3 with 5.5 mM Glucose) and counted. For mitochondrial membrane potential and oxygen consumption, we used protocols previously described^17,30,40,82^.

For mitochondrial membrane potential, 5 x10^7^ parasites was added to a cuvette previously prepared with intracellular buffer (125 mM sucrose, 65 mM KCl, 1 mM MgCl_2_ and 2.5 mM KH_2_PO_4_ in 10 mM HEPES-KOH buffer pH 7.2) containing 1 mM succinate, used as mitochondrial substrate. Safranin *O* (2.5 µM) was used as the fluorescent probe, and its fluorescence was measured in a Hitachi F-7000 fluorescence spectrophotometer. Digitonin (30 µM) was used to selectively permeabilize the plasma membrane, followed by 10 µM ADP, to stimulate ATP synthesis thus causing a depolarization of the mitochondrial membrane. Addition of 1 µg/mL of oligomycin would repolarize the membrane, and lastly addition of 5 µM FCCP depolarized the mitochondrial membrane.

The mitochondrial oxygen consumption rate was obtained using an Oroboros O2K respirometer (https://www.oroboros.at/). 10^8^ parasite was added to one of the chambers containing intracellular buffer and succinate. Parasites were permeabilized with digitonin, and the basal mCOR was measured. Addition of 200 µM ADP would stimulate the mOCR. Lastly 1.5 µM of oligomycin was added to obtain the minimal mOCR.

### Subcellular fractionation

The protocol for subcellular fractionation and mitochondrial membrane isolation was adapted from the one previously used for the purification of the plant like vacuolar compartment or acidocalcisomes^83,84^. Mutant parasites, either tagged with 3HA (*TgCoq5-3HA)* or with the modified biotin ligase turbo ID (*TgCoq5-TID*) were used to infect 10 T75 flasks per cell line (x10^9^ parasites in total). When using the *TgCoq5-TID* or the *FBXO14-TID* mutant, the growth media was supplement with 50 µM biotin for 2 h (TgCoq5-TID) or 1 h (FBXO14-TID) prior to collection. The pellet from the last wash was mixed with silicon carbide and grinding followed for approximately 2 min using an ice-cold mortar and pestle. When approximately 90% of the parasites were lysed, the mixture was transferred to an open top tube with lysis buffer (125 mM sucrose, 50 mM KCl, 4 mM MgCl_2_, 0.5 mM EDTA, 20 mM HEPES-KOH pH7.2, mini complete protease inhibitor, 12 µg/mL DNase, 12 µg/mL RNase, and 8 µg/mL nocodazole). Differential centrifugation speeds and times were shown in **Fig. S4B**. 500 µL of each soluble fractions were collected, and the pellets were resuspended in intracellular buffer. Aliquots from different fractions were then subjected to mitochondrial activity assay and western blots. The P2 fractions (higher mitochondrial activity) were used for blue native PAGE, 2D BN/SDS-PAGE, and for immunoprecipitation with streptavidin beads.

### Blue native and 2D BN/SDS-PAGE

The P2 fractions from the subcellular fractionation was solubilized in mitochondrial lysis buffer containing 2% dodecyl maltoside, 250 mM sucrose, 20 mM Tris HCl pH 8.0, 2 mM EDTA, and 750 mM amin-n-caproic acid as previously described^85^. 100 µg of solubilized protein or 10^8^ parasite lysate was loaded onto the first dimension BNPAGE gel and solubilized bovine heart mitochondria^41,86^ was loaded in the gel as molecular weight marker for mitochondrial complexes. At the end of the run, the lane containing the bovine heart mitochondria was cut off and stained with Coomassie. The lane containing the solubilized P2 fraction or parasite lysate was cut and denatured with 1X Laemmli buffer with 100 mM DTT and heated for 20 s in the microwave. After the gel lane was cooled to RT, it was transferred and run in a SDS PAGE gel (second dimension). The protein gel was then transferred onto a PVDF membrane at 25 V, 4°C overnight. The membrane was fixed with acetic acid and de-stained with methanol as previously described^85^. The membrane was then probed with the antibodies mentioned in the western blot section. Proteins were then imaged with a chemiluminescence using ChemiDoc. Each BNPAGE and BN/SDS-PAGE were performed at least twice with the representative images shown in the figures.

### Succinate cytochrome c reductase activity assay

The assay was carried out using a previously published protocol^87^. 2 µL of each fraction was used for each technical replicate (3 in total). After the addition of reaction buffer (0.3 mM succinate, 0.3 mM KCN, 10 mM HEPES) containing succinate as a substrate, the reaction started after the addition of cytochrome *c* (0.1 mM final). The absorbance of reduced cytochrome c was measured at 550 nm using a Biotek synergy H1 plate reader (Agilent, H1MFDG) for 10 min while shaking at 30°C. The activity of each fraction was then normalized based on their protein concentration measured by BCA.

### Pulldown of biotinylated proteins

Fractions were lysed in RIPA buffer (150 mM NaCl, 0.1 % SDS, 0.5 % sodium deoxycholate and 1% NP-40 in 50 mM HEPES pH 7.5 buffer) supplemented with mini complete protease inhibitor, and soluble protein concentrations were measured using BCA. 100 µg of proteins were used as input and bound to streptavidin magnetic beads overnight at 4°C. The experiment was carried as previously described^42^. In brief, the beads were washed with RIPA buffer, 1 M KCl, 0.1 M Na_2_CO_3_, 2 mM urea, and then twice with RIPA buffer. Proteins were eluted with 20 mM Biotin in 1% SDS and 25 mM Tris pH 8 by heating at 75°C for 30 mins. Elutes were sent for mass spectrometry analysis to identify enriched peptides.

### Reciprocal co-immunoprecipitation and identification of TgCoqFAD interactors

For reciprocal co-IPs, IPs were carried out as described previously^32^ with modifications. Protein-G magnetic beads (Thermo Fisher, 88847) were incubated with mouse αTgCoq3 or guinea pig αTgCoq5 in coupling buffer (10 mM NaH_2_PO_4_, 150 mM NaCl pH 7.2). Antibodies were crosslinked to the beads using 220 mM dimethyl pimelimidate (DMP, Sigma, D8388) in 200 mM sodium borate buffer (pH 9.0), and the reaction was quenched with 200 mM ethanolamine (pH 8.5). Parasites were lysed in buffer containing 1% digitonin, 20 mM HEPES (pH 7.4), 200 mM NaCl, and 2 mM MgCl₂, supplemented with protease inhibitor cocktail (Roche), for 1 h on ice. Lysates were clarified by centrifugation at 21,000 x g for 15 min at 4°C. An aliquot of the clarified lysates was saved as input and the remainder was incubated overnight at 4°C with αHA magnetic beads (Thermo Fisher, 88836) or with αTgCoq3- or αTgCoq5-coupled beads. Beads were washed twice with buffer containing 0.05% digitonin, 20 mM HEPES (pH 7.4), and 200 mM NaCl, followed by one wash with the same buffer containing 500 mM NaCl, and a final wash with 200 mM NaCl. Bound proteins were eluted by boiling in sample buffer for western blot analysis. For identification of TgCoqFAD interactors, TgCoqFAD-3HA and TgCoq1-3HA (control) were subjected to subcellular fractionation to enrich mitochondrial proteins. To preserve complex integrity, 10 µM UQ6 was included in the lysis buffer. P2 fractions were solubilized in buffer containing 1% digitonin, 20 mM HEPES (pH 7.4), 200 mM NaCl, and 10 µM UQ6, supplemented with protease inhibitors, and incubated for 2 h at 4 °C. Lysates were clarified by centrifugation at 21,000 × g for 30 min at 4 °C. Clarified lysates (200 µg, as determined by BCA assay) were incubated with αHA magnetic beads and washed as described above. Beads were additionally washed with PBS (pH 7.4) prior to on-bead digestion and mass spectrometry analysis.

### Protein identification using LC-MS/MS

Samples were sent to the Proteomics and Metabolomics Facility, Nebraska Center for Biotechnology, University of Nebraska-Lincoln and analyzed for mass spectrometry (MS) as previously described^88^. Briefly, an aliquot of 37.5 µL of sample was added to 12.5 µL of 4X reducing NuPAGE LDS gel (Thermo Fisher Scientific, Waltham, MA) sample buffer at 5 mM dithiothreitol (DTT) and incubated at 95°C for 10 min. The samples were loaded and run on a Bolt 12% Bis-Tris-Plus gel (Thermo Fisher Scientific) in MES SDS running buffer to clean-up the samples and concentrate the proteins into the top of the gel. The gel was then fixed in methanol:acetic acid:water (40:10:50) and stained with Colloidal Coomassie blue G-250. The gel containing proteins was excised and destained in 50% acetonitrile (ACN), 50 mM ammonium bicarbonate. The proteins were reduced in 100 mM ammonium bicarbonate with 10 mM DTT. The reducing buffer was removed, and proteins were alkylated with iodoacetamide 10 mM. Proteins were digested with 250 ng of trypsin overnight at 37°C. Peptides were extracted from the gel pieces, dried down, and re-dissolved in 5% acetonitrile, 0.2% formic acid. Each digest was run by nanoLC-MS/MS using a 2 h gradient on a Waters CSH 0.075 mm x 250 mm C18 column (Waters Corp, Milford, MA) feeding into a Thermo Orbitrap Eclipse mass spectrometer.

All MS/MS samples were analyzed using Mascot (Matrix Science, London, UK; version 2.7). Mascot was set up to search the cRAP_20150130.fasta (125 entries); and ToxoDB-59_TgondiiGT1_AnnotatedProteins_20221003 (8,460 sequences) assuming the digestion enzyme trypsin. Mascot was searched with a fragment ion mass tolerance of 0.060 Da and a parent ion tolerance of 10.0 PPM. Carbamidomethyl of cysteine was set as a fixed modification. Deamidated of asparagine and glutamine, oxidation of methionine was specified in Mascot as variable modifications. Scaffold (version Scaffold_4.8.9; Proteome Software Inc., Portland, OR) was used to validate MS/MS-based peptide and protein identifications.

Hits were identified with 95% protein threshold and peptide threshold, and the minimum number of peptides detected was set at 2. Statistical analysis was done using Fisher’s exact test with Benjamini-Hochberg multiple test correction to identify significantly enriched proteins. To obtain a numeric fold-change value, peptides with values of 0 were adjusted to 0.0001.

### Analysis of quinone content

Parasite pellets were resuspended in 0.4 ml of 95% (vol/vol) ethanol, transferred to a 5-ml pyrex tissue grinder, and homogenized for 1 min. Homogenates were transferred to a 10-ml pyrex tube containing ∼1 ml of 0.1 mm zirconia/silica beads (Biospec Products # 11079101z). The tissue grinder was rinsed twice with 0.3 ml of 95% (vol/vol) ethanol, and washes were combined to the original extracts. Samples were vortexed vigorously for 3 min and then heated at 70°C for 10 min. Samples were vortexed again for 3 min while cooling down and then mixed with 0.5 ml of water. The extracts were then partitioned three times with 5 ml hexane. Hexane layers were combined, evaporated to dryness with gaseous nitrogen, and resuspended in 0.3 ml of methanol:dichloromethane (10:1). Extracts were analyzed by HPLC on a 5 µM Supelco Discovery C-18 column (250 x 4.6 mm, SigmaAldrich) thermostated at 30°C and developed isocratically at a flow rate of 1 ml/min with methanol:hexane (95:5; vol/vol). DMQ_7_ (10.5 min), UQ_7_ (11.3 min) and UQ10 (32.2 min) were detected spectrophotometrically at 275 nm and quantified according to external calibration curves. The corresponding quinol versions were fully re-oxidized during the heating step and were quantified as part of the quinone pool. UQ_10_ standard was from Sigma-Aldrich; UQ_7_ standard was extracted from *Candida utilis* and purified via reverse-phase chromatography. DMQ_7_ was HPLC-purified from *iΔTgCoqFAD* cell extracts (+ATc) and its identity was verified using a hybrid quadrupole orthogonal time of flight spectrometer (Agilent technologies) using the same conditions as those described in Latimer et al (2021)^45^. Ammonium adduct of DMQ_7_ was detected at m/z 782.65.

### Computational modelling, virtual screening and docking

A model of the full length mature (i.e. devoid of the first 149 amino acids that are cleaved upon mitochondrial targeting) TqCoqFAD (Uniprot accession no. A0A139Y5I3) was modelled along with FAD as co-factor using AlphaFold 3^89^. The best ranked model was further refined through GalaxyRefine server^90^ which significantly improved the quality of the model further and the final model had the Molprobity score of 1.568, clash score of 11.3, 0.1% poor rotamer and 99.1% residues as Ramachandran favored.

Three potential druggable pockets proximal to the FAD molecule onto the refined TgCoqFAD model were predicted by FTSite^48^. Using a GRID box spanning these potential druggable pockets, virtual screening (VS) was performed with DrugBank 6.0 (approved) library^49^ using FRED implemented in OEDocking 4.3.2 suite (OpenEye, Cadence Molecular Sciences, Inc., Santa Fe, NM) at its default setting. From the top 100 hits, molecules with total polar surface area (TPSA)>80 were excluded and the remaining ones were blindly docked to the TqCoqFAD model using AutoDock Vina version 1.1.2^91^ with exhaustiveness of 16. The hits preferentially going to the desired sites were subjected to focused docking using GOLD 5.3 suite (CCDC, Cambridge). Finally, the best 3 molecules were chosen for experimental testing, considering pose and docking scores used by Vina and GOLD. All ligand molecules for docking were obtained from PubChem.

For Figure 7B, 2D ligand interaction diagrams were generated using PoseView through Protein Plus server^92^. Black dotted lines indicate hydrogen bonding; green dotted lines indicate p-p stacking whilst green contour lines indicate hydrophobic and van der Waals interactions.

### Small molecule inhibition and host cell viability

EC_50_ of the compounds were obtained by following the protocol as previously described^81^. In brief, hTERT cells were seeded onto black 96 well plates and cultured for two days prior to the addition of 4,000 *TatiΔKu80* RFP or *iΔTgCoqFAD-3HA* RFP parasites per well and various concentrations of the small molecules. Fluorescence of RFP was measured on the plate reader and the EC_50_ values were calculated using GraphPad Prism. hTERT cells viability was assayed using Alamar Blue as described^81^ using 10X EC_50_ of the compounds against *TatiΔKu80*. The EC_50_s were calculated using GraphPad Prism software.

### *In vitro* bradyzoite viability assay

The assay was carried out as described previously^93^. In brief, confluent HFF monolayers were preincubated with 3 µM compound 1, 4-[2-(4-fluorophenyl)-5-(1-methylpiperidine-4-yl)-1H-pyrrol-3-yl] pyridine, for 2 hours before infection with *ME49ΔKu80*. After 24 h, the medium was replaced, and cells were cultured at 37°C under ambient CO_2_ conditions for 3 days to allow differentiation. Differentiated bradyzoites were then treated with drugs at 3X EC_50_ for 4 days. Bradyzoites were liberated by acid/pepsin digestion (170 mM NaCl, 60 mM HCl, and 0.1 mg/mL pepsin) for 10 min in a 37°C water bath and neutralized with 94 mM Na_2_CO_3_. A total of 10,000 bradyzoites were plated per well in a fresh confluent HFF six-well plate and allowed to grow in a 37°C incubator with 5% CO_2_ for 16 days. Plates were then fixed with 100% ethanol and stained with crystal violet as described in the plaque assay section. Plaques were counted using a light microscope (Nikon Eclipse TS100) with a 4X objective.

### Statistical analysis

Experimental data were expressed as the mean with standard error of the mean (SEM) from at least three biological repeats unless indicated otherwise. Statistical analyses were performed using GraphPad PRISM (version 10). Student’s t-test, one-way ANOVA, and two-way ANOVA tests were used when appropriate and indicated in the figure legends. Significance was considered when *p*-value was less than 0.05.

