## Supplementary Figures and Legends for "Evolutionary Remodeling of Ubiquinone Biosynthesis in *Toxoplasma gondii* Reveals an Essential Bi-functional Monooxygenase"

**Supplementary Figure 1. Ubiquinone biosynthesis pathway from mammals and bacteria**

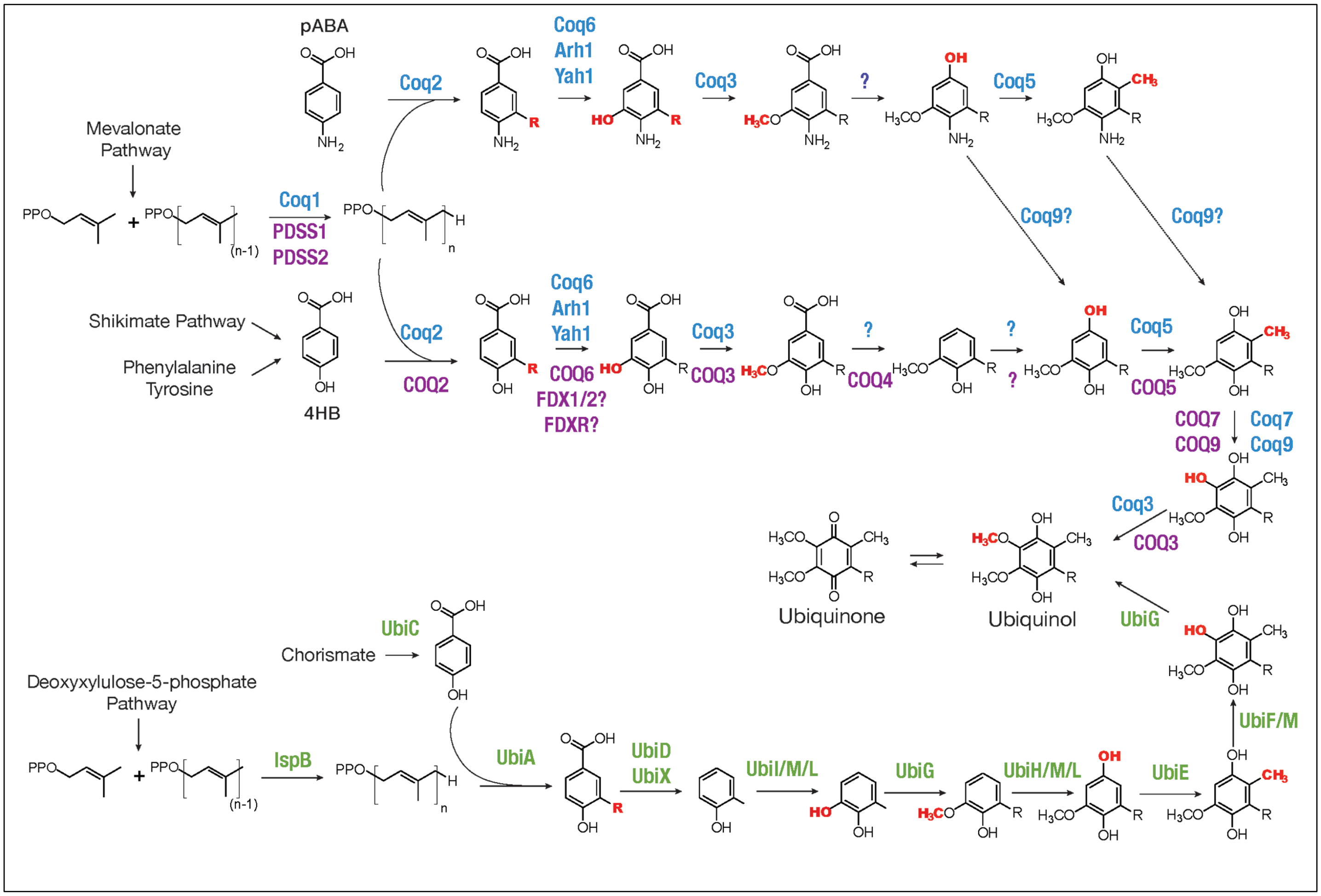


Reactions involved in UQ biosynthesis in yeast (enzyme names in dark blue), mammals (enzyme names in purple), and bacteria (enzyme names in green). Proteins that have not yet been identified are indicated with a question mark. 4HB, 4-hydroxybenzoic acid; pABA, para-aminobenzoic acid. In mammals, both phenylalanine and tyrosine can be used for 4HB synthesis, whereas yeast utilizes only tyrosine. UbiM and UbiL are present only in certain bacterial species.

**Supplementary Figure 2. Regulation of the expression of TgCoq3 and TgCoq5**

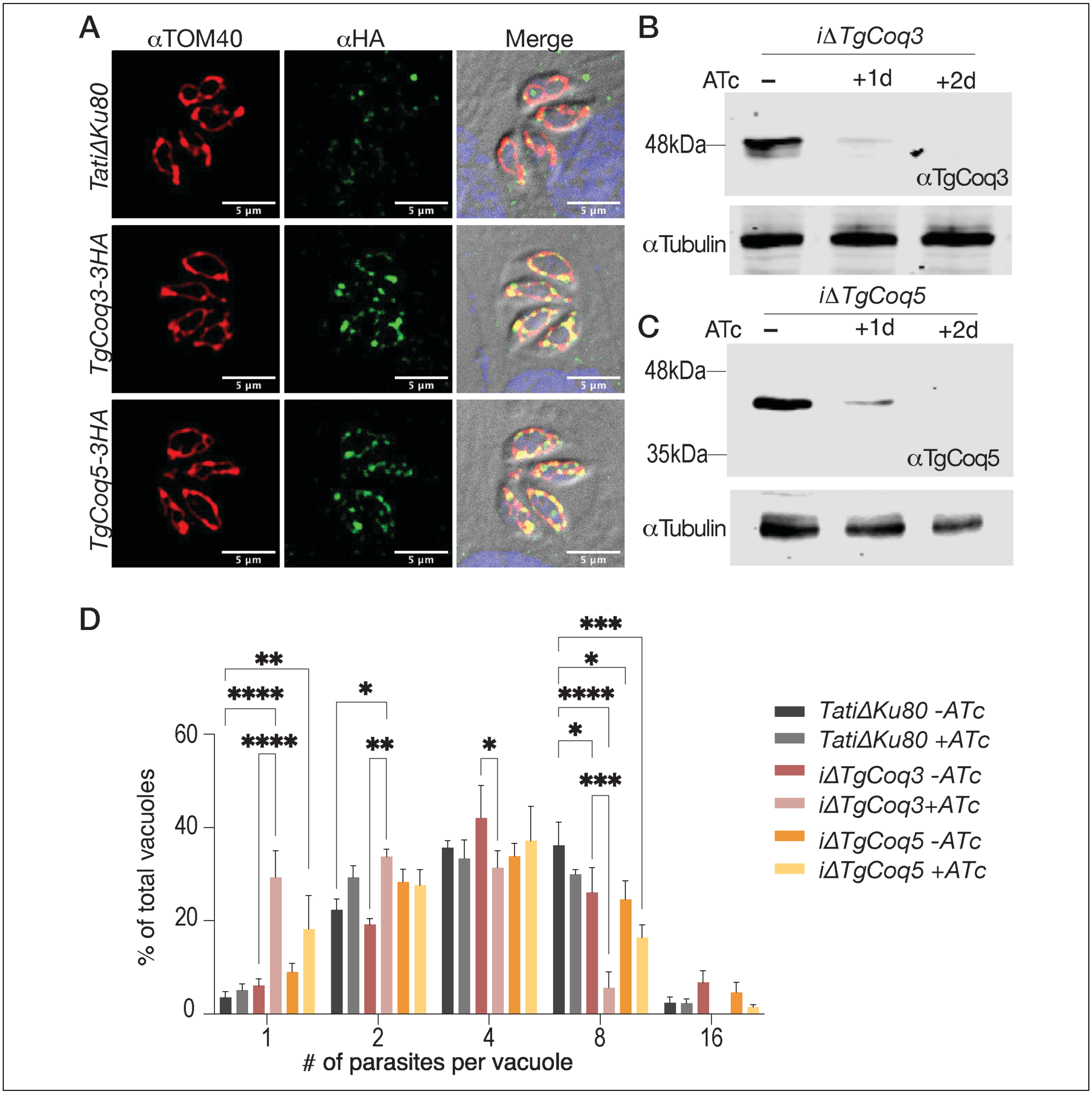


(A) IFA of C-terminally tagged HA cell lines. αTOM40 was used to label mitochondria and TgCoq3 and TgCoq5 were probed with αHA. *TatiΔku80* was used as negative controls. (B) Western blot of total lysates of the *iΔTgCoq3* mutant incubated with ATc for the indicated times. Mouse αTgCoq3 was used to probe for TgCoq3, and αTubulin was used as loading control. (C) Western blot of proteins obtained from lysates of the *iΔTgCoq5* mutant grown with ATc for the indicated times. Guinea pig αTgCoq5 was used to detect TgCoq5, and αTubulin was used as loading control. (D) Number of parasitophorous vacuoles counted in host cells infected with the indicated cell lines for 20 hours ± ATc for three days.

Data was obtained from three biological repeats. Data was analyzed with Two-way ANOVA, and the bars represent SEM. $*p \leq 0.05$, $**p \leq 0.01$, $***p \leq 0.001$.

**Supplementary Figure 3. Generation and purification of αTgCoq3 and αTgCoq5 antibodies**

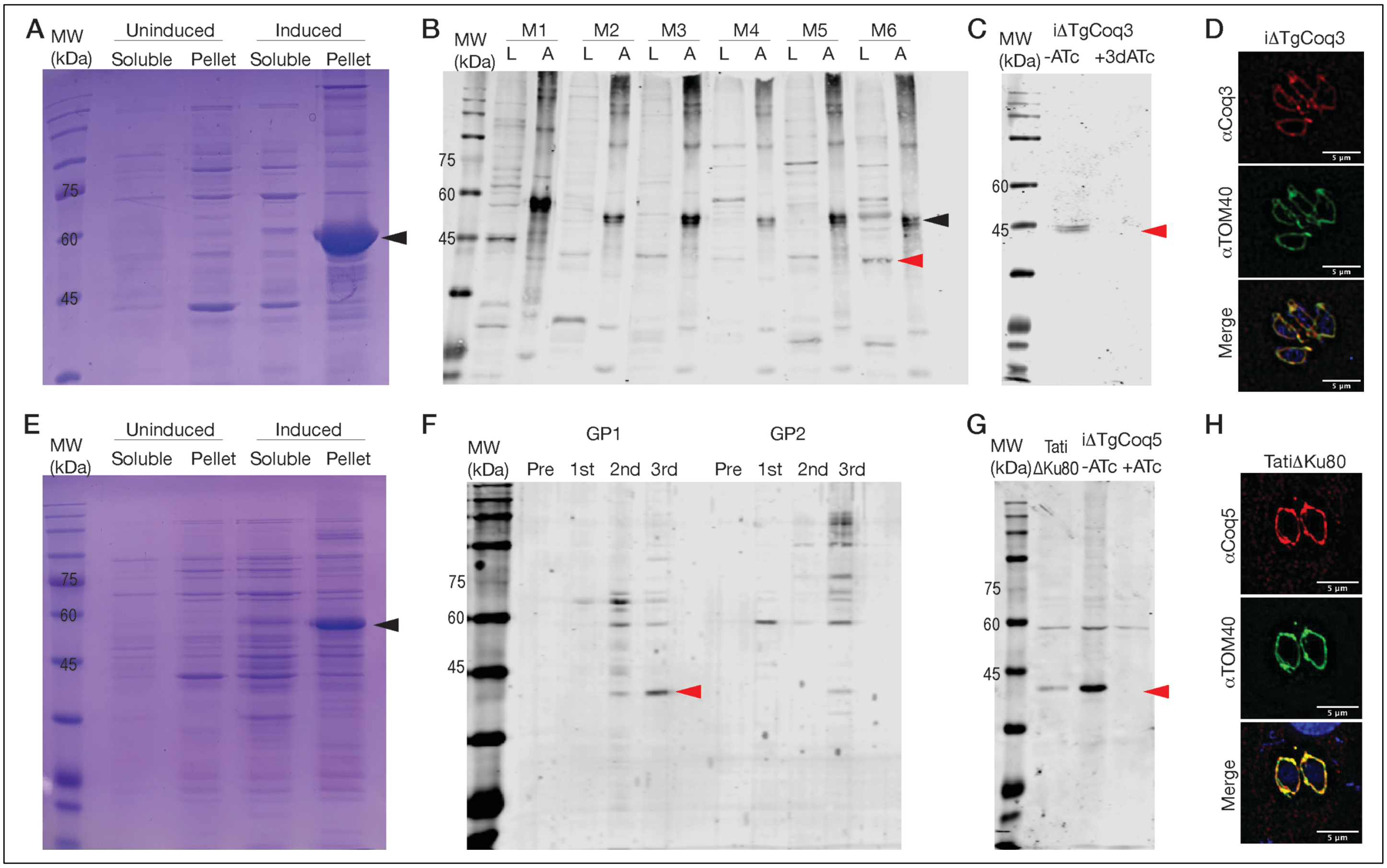


(A) Coomassie gel showing TgCoq3 induction. Black arrowhead points to TgCoq3. MW: molecular weight marker. (B) Western blot probed with serum collected from 6 mice (M1-M6) one week after the last boost. L: *TatiΔku80* lysate, A: diluted antigen. Black arrowhead: TgCoq3 antigen, red arrowhead: mature TgCoq3 protein. (C) Western blot using lysates from *iΔTgCoq3* either not preincubated with ATc or preincubated with ATc for 3 days. Membrane probed with affinity purified αTgCoq3 from M3. (D) IFA showing αTgCoq3 colocalization with the mitochondrial marker TOM40. (E) Coomassie gel showing TgCoq5 induction. Black arrowhead: TgCoq5. (F) Western blot using *TatiΔku80* lysate probed with serums from 2 guinea pigs (GP1 and GP2). Pre: pre-immunization serum. 1st: after first boost, 2nd: after second boost, and 3rd: after third boost. Red arrowhead: mature (with mitochondrial targeting peptide cleaved) TgCoq5. (G) Western blot using lysates from *TatiΔku80* and *iΔTgCoq5* with or without 3 days of ATc treatment. Red arrowhead: mature TgCoq5. (H) IFA showing αTgCoq5 colocalization with the mitochondrial marker TOM40.

**Supplementary Figure 4. Mitochondrial enrichment by subcellular fractionation**

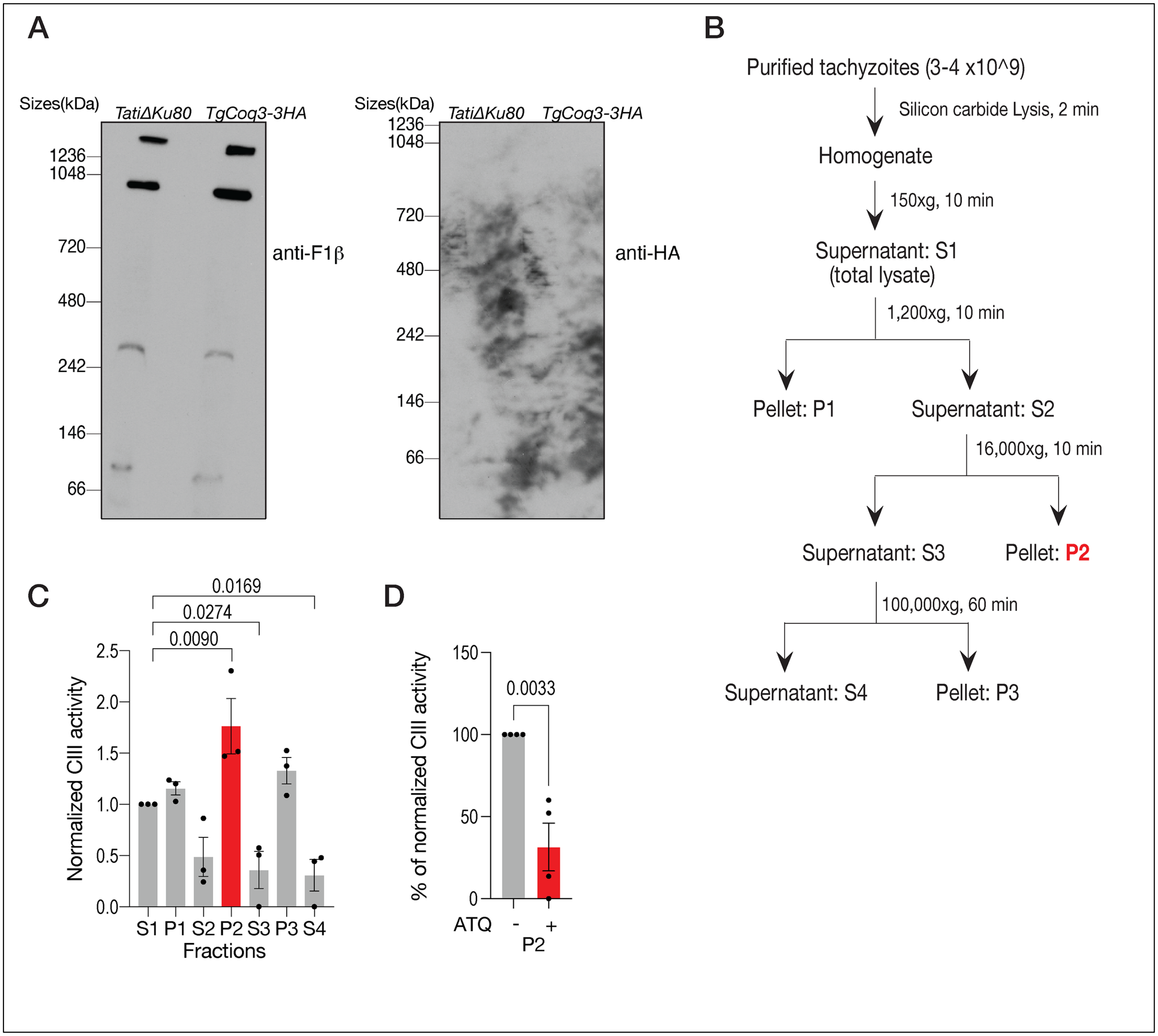


(A) BNPAGE using 2x10$^7$ *TatiΔku80* or *TgCoq3-3HA* parasites lysed with 2% digitonin. NativeMark was used as molecular weight marker. Left, membrane probed with αF1β (ATP synthase subunit). Right, same membrane probed with αHA. (B) Schematics of the subcellular fractionation protocol used to obtain enriched mitochondrial fractions. (C) Succinate cytochrome *c* reductase activity using the fractions obtained from three subcellular fractionations. Each point represents a biological sample, the bars represent SEM. One-way ANOVA test was used, n=3. (D) Cytochrome c reduction assay with P2 fractions either incubated with vehicle control (DMSO) or 10 nM atovaquone (ATQ) for 10 minutes before beginning the assay. Cytochrome c reduction activity was normalized to DMSO control. Each point represents a biological replicate, and the bar represent SEM. Student's t-test was used, n=4. Significance was considered when p-values are less than 0.05.

**Supplementary Figure 5. TgCoq4 and TgCoq8 localize to the mitochondrion and interact with TgCoq5.**

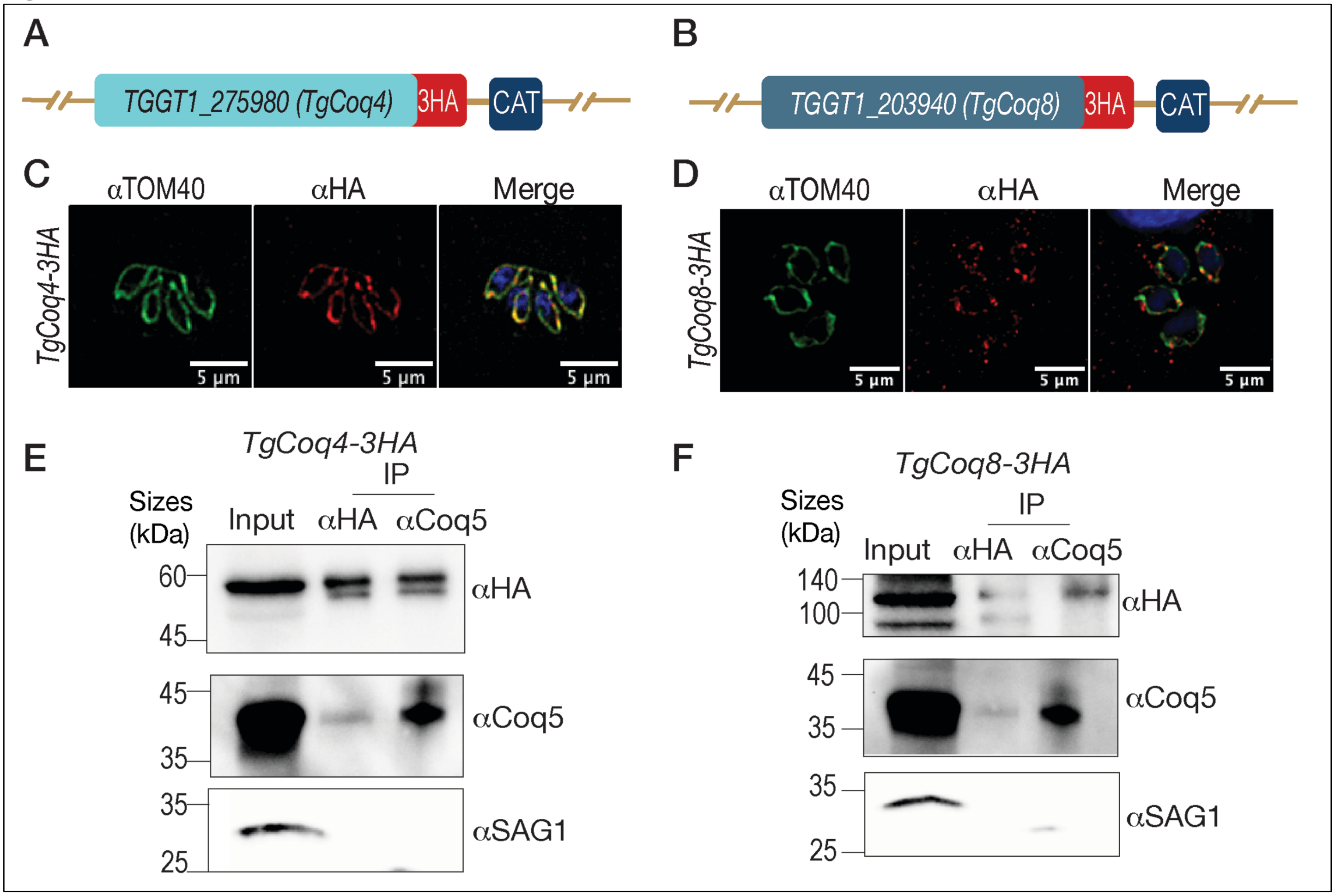


(A) Scheme showing tagged TgCoq4 with 3HA and the chloramphenicol acetyl transferase (CAT) gene used for selection with chloramphenicol. (B) Scheme showing tagged TgCoq8 with 3HA and CAT drug resistant cassette. (C) IFAs using αHA to probe for *TgCoq4-3HA*. TOM40 was the mitochondrial marker. (D) IFAs using αHA to probe for *TgCoq8-3HA*. TOM40 was used as the mitochondrial marker. (E) Reciprocal co-IPs showing interactions between TgCoq4 and TgCoq5 using αHA- and αTgCoq5-coupled beads. TgCoq4-HA was detected with αHA, while TgCoq5 was detected with αTgCoq5 and SAG1 served as negative control. (F) Reciprocal co-IPs showing interactions between TgCoq8 and TgCoq5 using αHA and αTgCoq5-coupled beads. TgCoq8-αHA, while TgCoq5 was detected with αTgCoq5, and SAG1 was used as control. The reciprocal co-IPs were repeated three times.

**Supplementary Figure 6. TgCoq5-TID requires ≥2 h of biotin labeling to detect mitochondrial proteins, and whole-cell lysate immunoprecipitation does not enrich for mitochondrial components.**

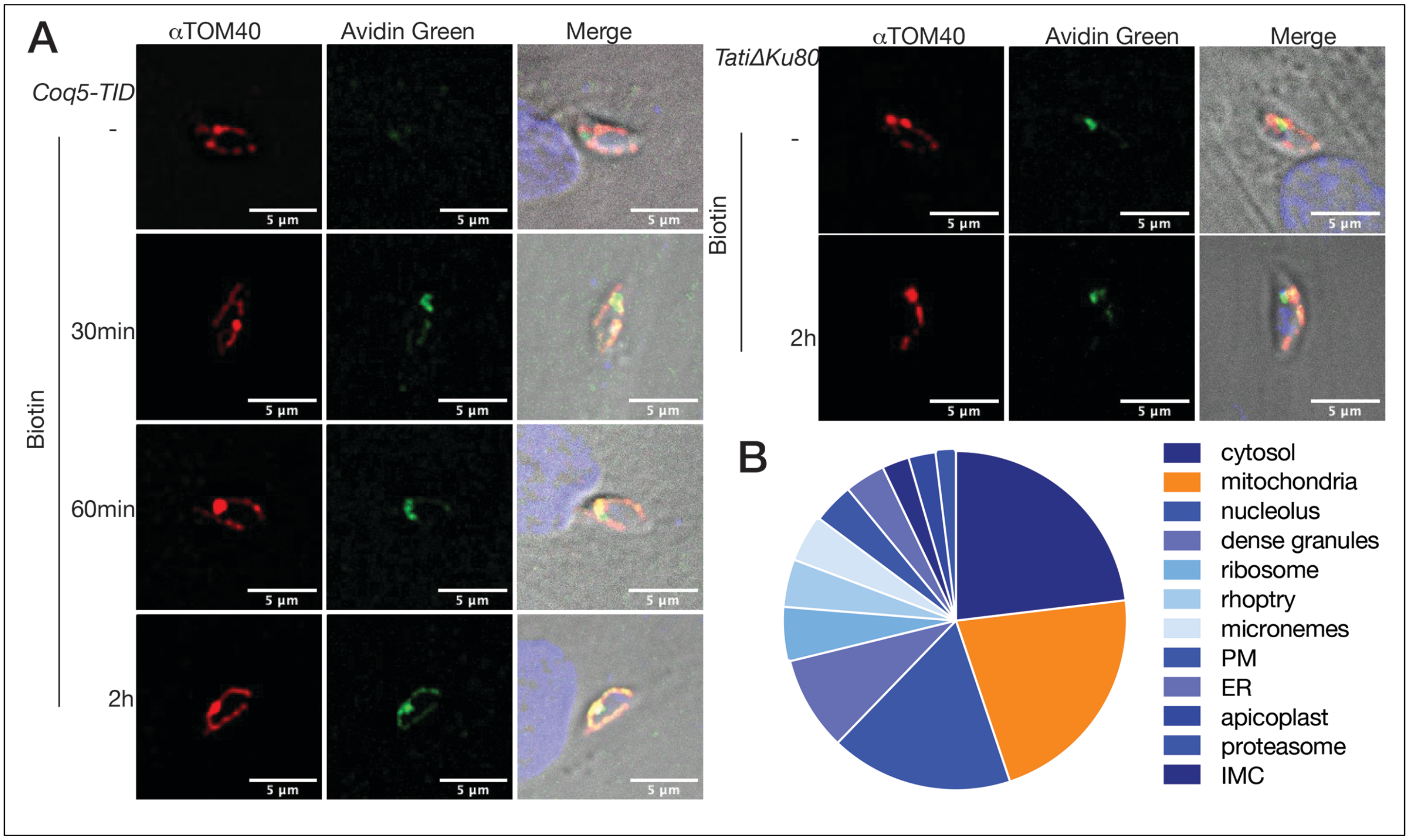


(A) IFA of TgCoq5-TID cell line incubated with biotin for various lengths of time. Biotinylated proteins were probed with avidin green and TOM40 was used as mitochondrial marker. (B) Pie chart of the protein localization predicted by LOPIT[1] of peptides enriched in *TgCoq5-TID/TatiΔku80* whole cell lysates after streptavidin IP.

**Supplementary Figure 7. Characterization of candidate hypothetical proteins listed in Table 1**

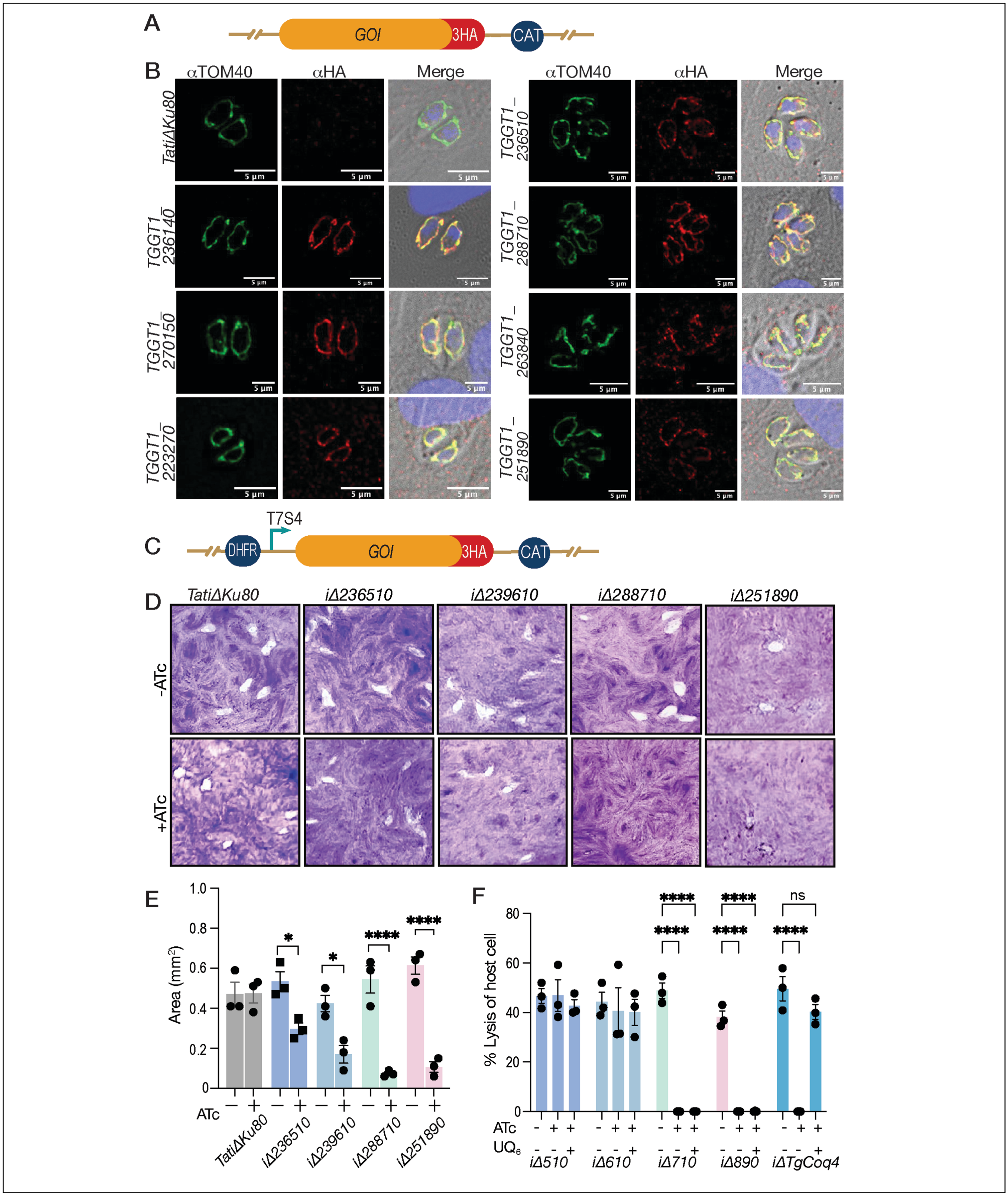


(A) Schematics for tagging each gene of interest (GOI) with 3xHA at their C-terminus. CAT: chloramphenicol acetyl transferase cassette for selection. (B) IFA of C-terminally HA tagged hypothetical proteins from Table 1. TOM40 was used as mitochondrial marker. (C) Scheme showing the plasmid structure used for generating conditional knockdowns of each GOIs. DHFR: dihydrofolate reductase used for selection with pyrimethamine. T7S4: tetracycline response element and SAG4 promoter. (D) Representative plaque assays of four of the conditional knockdown cell lines of the hypothetical proteins. Image taken at day 7. (E) Quantification of plaques formed by the mutants from part D. +ATc: parasites preincubated with ATc for 3 days prior to the assay. Data from three biological repeats and data was analyzed with Two-way ANOVA. Each point represents a biological replicate, and the bars represent SEM. ($*p \leq 0.05$, $**p \leq 0.01$, $***$ $p \leq 0.001$, $****$ $p \leq 0.0001$. (F) 96 well plate crystal violet assay measuring the percentage of lysed host cell by the same mutants shown in E plus *iΔTgCoq4*. Mutant parasites were either incubated without ATc, with ATc, or with ATc plus 10 μM $UQ_6$. Plate was fixed and stained at day 5. Percentage of lysis was calculated using host cell only control. Each point represents a biological replicate, n=3, and the bars represent SEM. $*p \leq 0.05$, $**p \leq 0.01$, $***$ $p \leq 0.001$, $****$ $p \leq 0.0001$.

**Supplementary Figure 8. Phylogenetic tree of TgCoqFAD orthologs and paralogs in diverse**

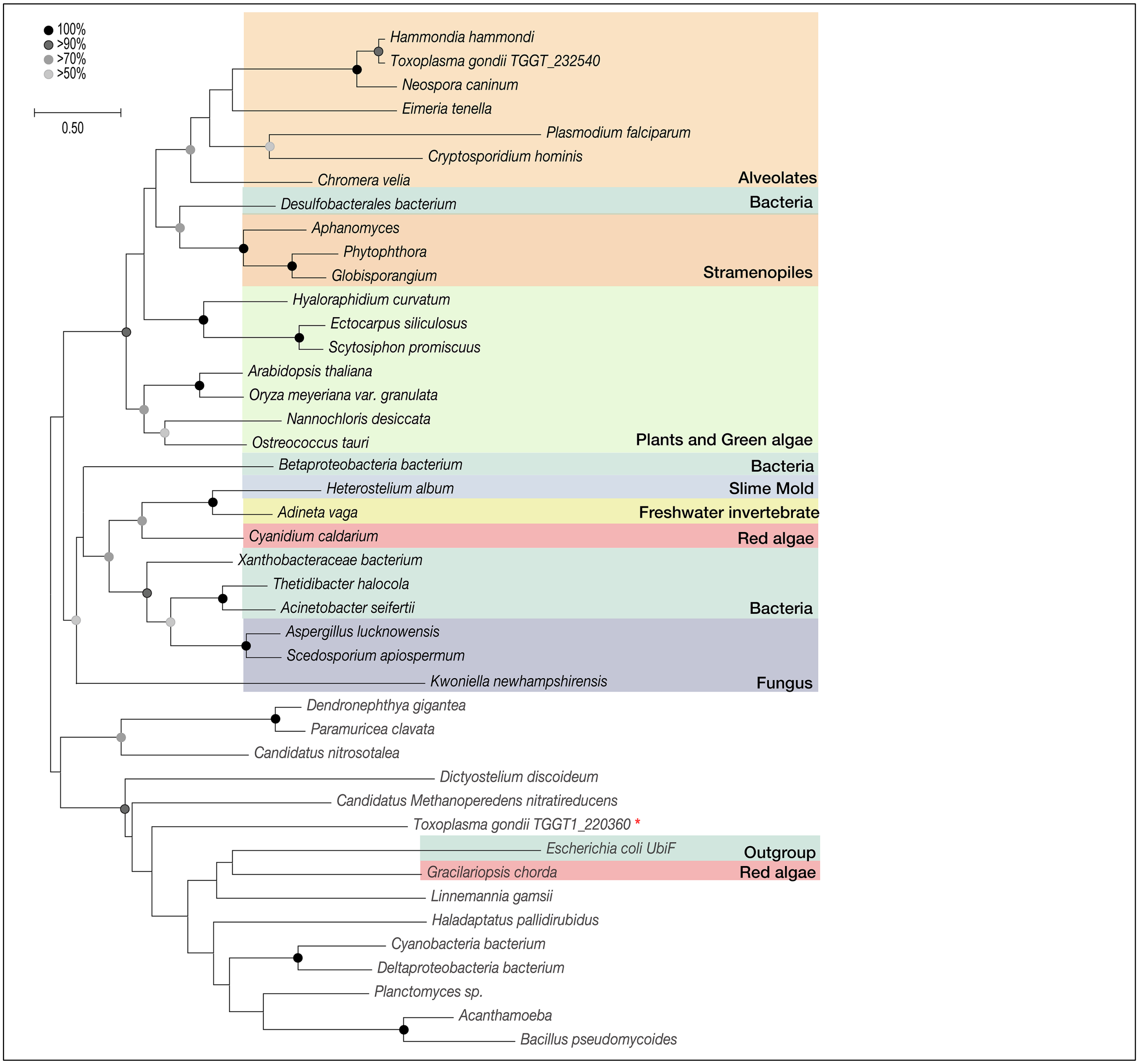


Extended phylogeny tree of TgCoqFAD orthologs and homologs in different organisms. Sequences used to generate this tree is listed in **Supplementary table 5**. *E. coli* UbiF was used as an outgroup. Red asterisk indicates the non-mitochondrial FAD-dependent monooxygenase in *T. gondii*.

**Supplementary Figure 9. Knockdown of TgCoqFAD causes buildup of an UQ intermediate and the growth defect cannot be bypassed with benzoic acid analogs**

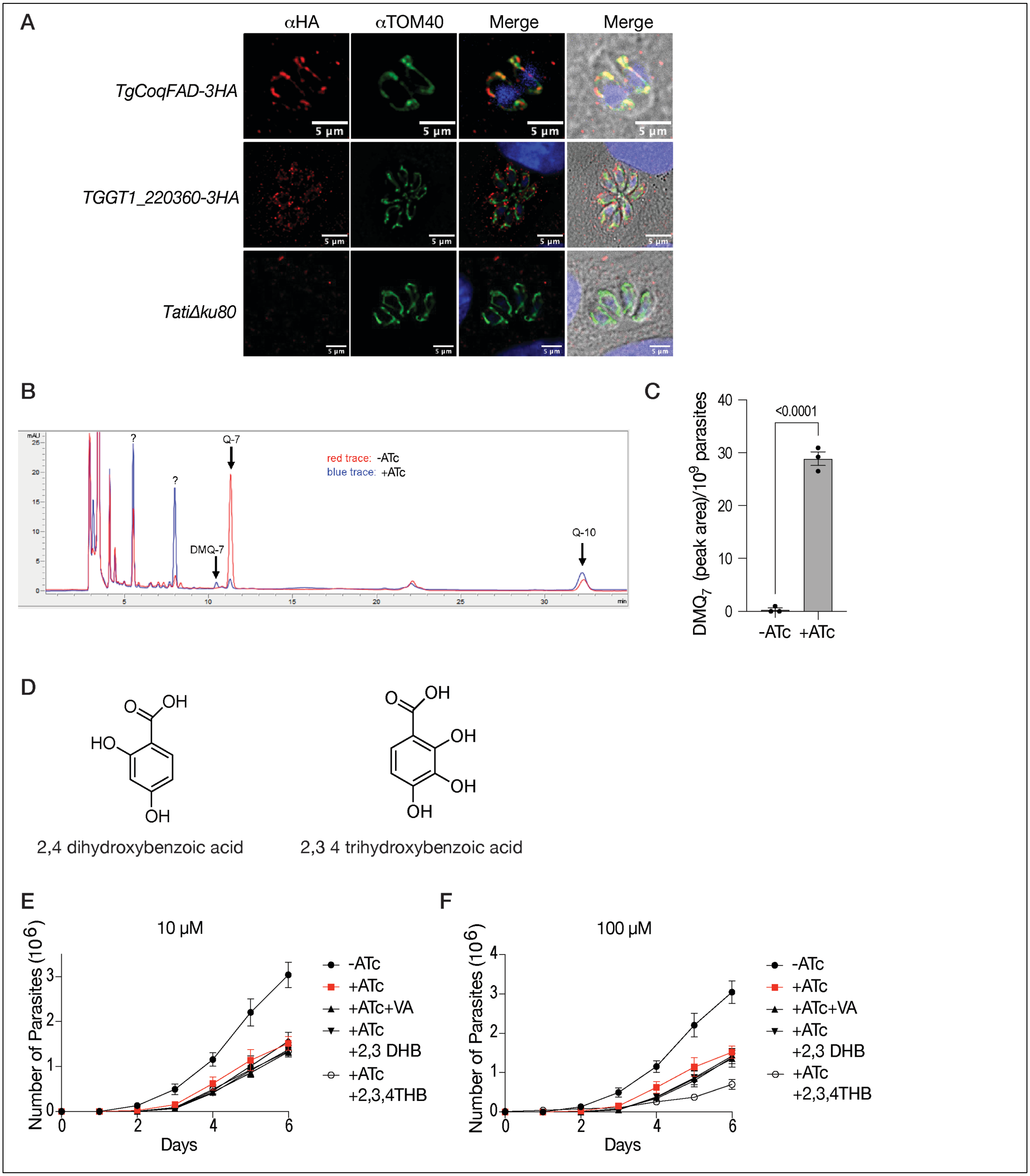


(A) IFA of *TgCoqFAD-3HA* and *TGGT1_220360-3HA* parasites. αTOM40 was used as a mitochondrial marker. (B) HPLC analysis of lipids extracted from *iΔTgCoqFAD* ± ATc for six days. The peaks corresponding to $UQ_7$, $UQ_{10}$, and demethoxy-ubiquinone$_7$ ($DMQ_7$) are indicated with an arrow. (C) Quantification of the peak area corresponding to the intermediate $DMQ_7$ from *iΔTgCoqFAD* ± ATc for six days. Data presented as mean with SEM from three biological replicates, and student's t test used for statistical significance. (D) Structures of hydroxybenzoic acid analogs used in E and F. (E) Growth of TgCoqFAD expressing RFP ± ATc and +ATc plus 10 μM of VA (Vanillic acid), 2,4 DHB (dihydroxybenzoic acid), or 2,3,4 THB (tri-hydroxybenzoic acid). Parasites were preincubated with ATc for three days prior to the infection. Data presented as mean with SEM from three biological replicates. (F) Growth assay similar as in E, except that 100 μM of the analogs were used.

**Supplementary Figure 10. TgCoqFAD knockdown does not affect the UQ synthesis complex formation. Expression of the *Arabidopsis thaliana* FMO in the mitochondrion of the *i*Δ*TgCoqFAD* mutant**

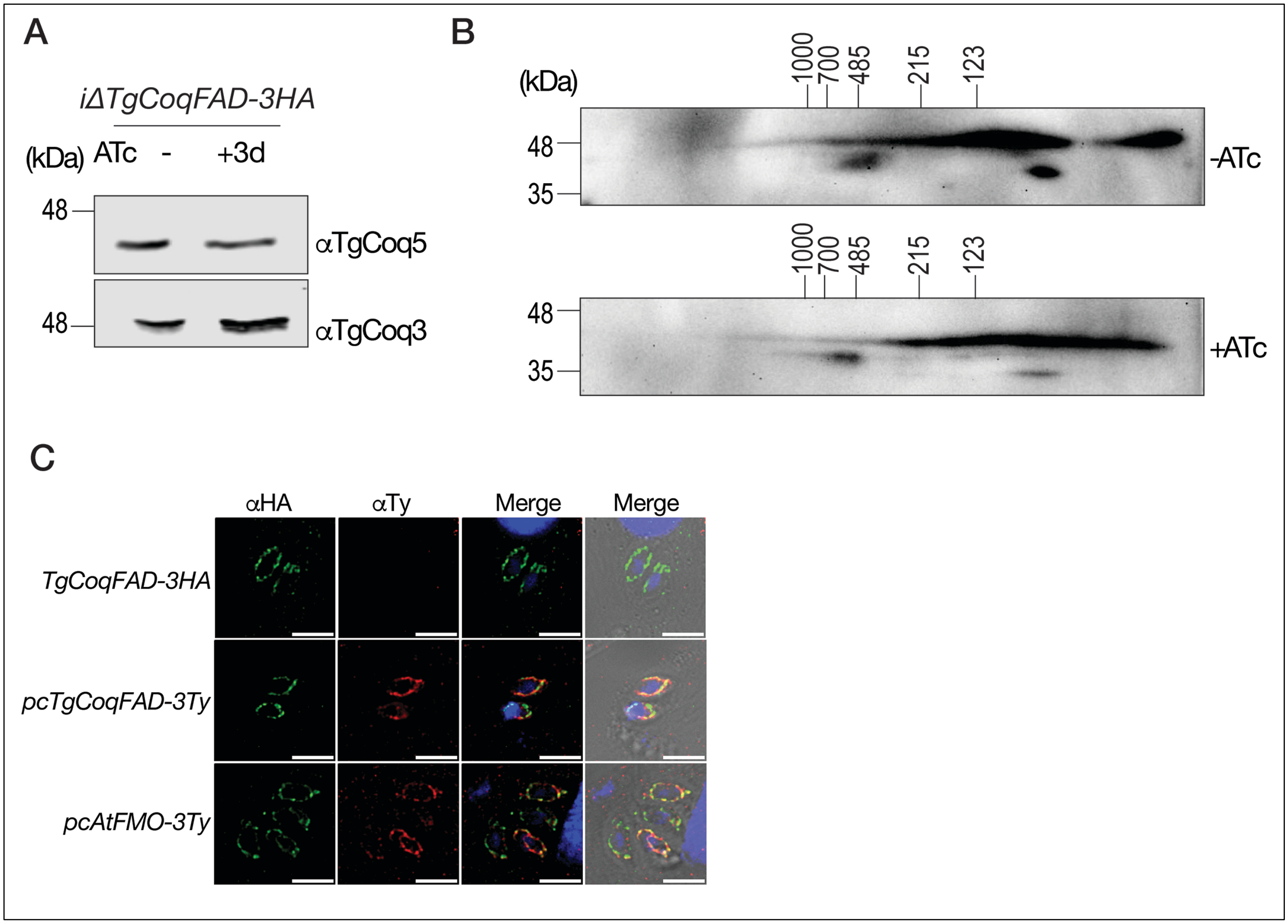


(A) Western blot using whole cell lysates from *i*Δ*TgCoqFAD* ± 3 days of ATc. Membrane from **Fig 5D** stripped and probed with αTgCoq3, and αTgCoq5. (B) 2DPAGE analysis using lysates from *i*Δ*TgCoqFAD* ± ATc for three days. UQ complex was visualized using αTgCoq5. (C) IFA showing the expression and mitochondrial localization of TgCoqFAD-3Ty and AtFMO-3Ty transiently transfected into *TgCoqFAD-3HA* cell line. αHA was used to probe for endogenous TgCoqFAD-3HA, αTy was used to probe for transiently expressed TgCoqFAD-3Ty and AtFMO-3Ty. Scale bars represent 5 µm.

**Supplementary Figure 11. Potential druggable pockets identified in TgCoqFAD**

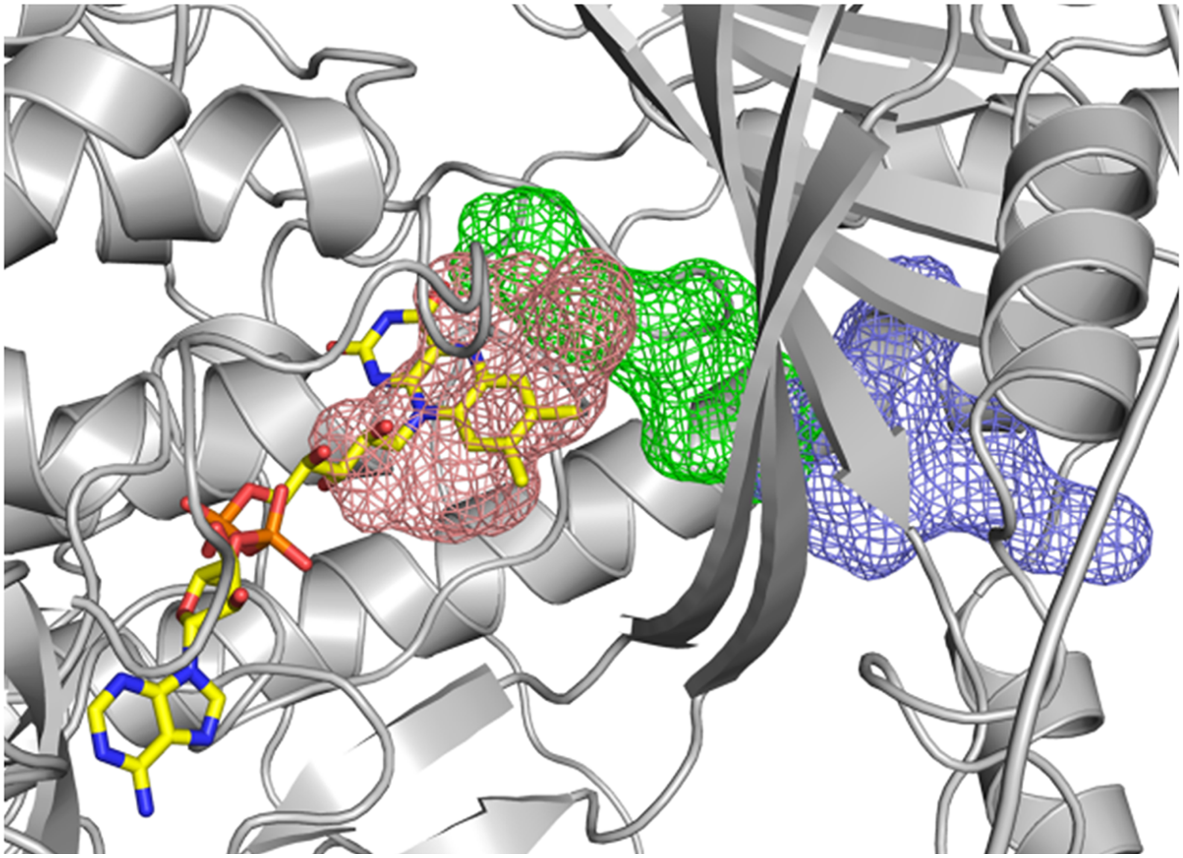

The protein is shown as a light grey cartoon, with the modeled FAD displayed as sticks. The three highest-ranked potential ligand-binding pockets, identified using FTSite, are shown as meshes in salmon, green, and pale blue. The figure was generated using PyMOL.

**Supplementary Figure 12. Overexpression of TgCoqFAD and validation of *in vitro* cyst formation.**

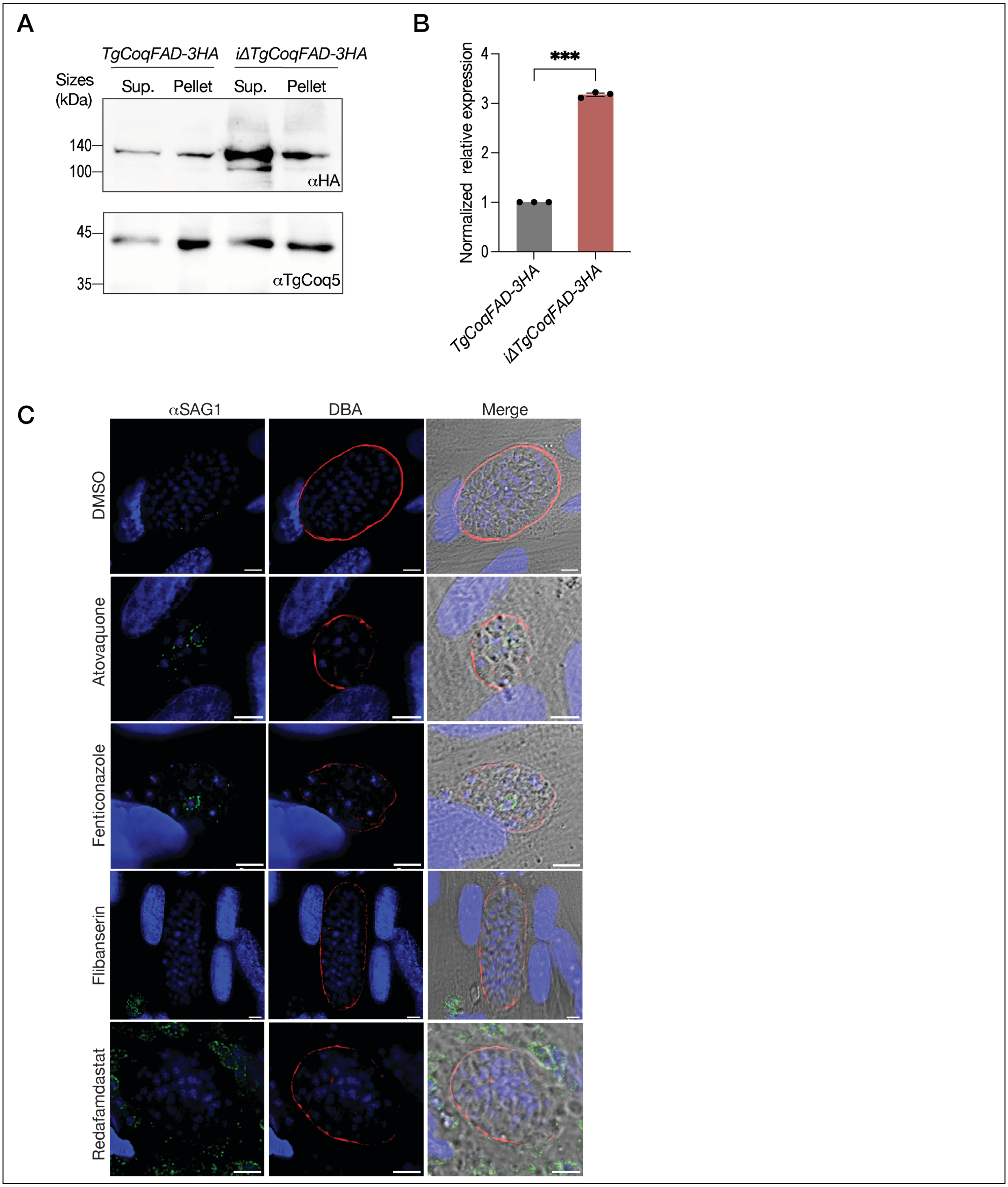


(A) Membrane (pellet) and soluble (supernatant) protein fractions from *TgCoqFAD-3HA* and *i*Δ*TgCoqFAD-3HA* mutants were prepared by freeze-thaw. Twenty µg of each fraction were loaded per lane. TgCoqFAD-3HA was detected with αHA; αTgCoq5 served as a loading control. (B) Quantification of total TgCoqFAD expression relative to total TgCoq5 expression normalized to *TgCoqFAD-3HA*. Data are from three biological replicates, shown as mean ± SEM; each dot represents one replicate. Statistical significance was assessed using Student's t-test, *** $p \leq 0.001$. (C) IFA of drug treated *in vitro* differentiated cysts were treated with 3X $EC_{50}$ of the compounds for 4 days (same condition as Figure 7D and 7E prior to bradyzoite liberation), and the cyst wall was stained with DBA while SAG1 is used to probe for tachyzoite surface antigen. Scale bars represent 5 μm.
