## Supplementary Tables for "Evolutionary Remodeling of Ubiquinone Biosynthesis in *Toxoplasma gondii* Reveals an Essential Bi-functional Monooxygenase"

### Supplementary Table 1. Predicted *T. gondii* UQ synthesis pathway proteins

| *T. gondii* ID | Annotation/ E-value | Yeast gene | Fitness/ Localization[1] | Reference |
|---|---|---|---|---|
| TGGT1_269430 | heptaprenyl diphosphate synthase COQ1/ 8e-44 | Coq1 | -3.66/ Mitochondria | 2 |
| TGGT1_259130 | 4-hydroxybenzoate polyprenyl transferase/ 5e-43 | Coq2 | -4.07/ Mitochondria | Predicted |
| TGGT1_266850 | 3-demethylubiquinone-9 3-O-methyltransferase/ 4e-22 | Coq3 | -4.49/ Mitochondria | This work |
| TGGT1_275980 | coenzyme q (ubiquinone) biosynthesis protein coq4 protein/ 2e-24 | Coq4 | -5.08/ Mitochondria | This work |
| TGGT1_295690 | ubiquinone/menaquinone biosynthesis methyltransferase subfamily protein/ 1e-41 | Coq5 | -3.62/ Mitochondria | This work |
| TGGT1_261030 | pyridine nucleotide-disulfide oxidoreductase domain-containing protein/ 6e-52 | Arh1 | -3.66/ Mitochondria | Predicted |
| TGGT1_240670 | adrenodoxin-type ferredoxin, putative/ 1e-40 | Yah1 | -2.45/ Mitochondria | Predicted |
| TGGT1_220360 | FAD binding domain containing protein/ 7e-31 | Coq6 | -0.9/ Plasma membrane | This work |
| No data | | Coq7,9,11 | | |
| TGGT1_203940 | ABC1 protein/ 2e-73 | Coq8 | -4.2/ Mitochondria | This work |
| TGGT1_233550 | polyketide cyclase/dehydrase/ 2e-08 | Coq10 | -5.54/ No LOPIT | Predicted |

The *Saccharomyces cerevisiae* UQ synthesis protein sequences were obtained from UniProt[3]. Candidate *T. gondii* homologs were identified by BLASTP[4] searches against the *T. gondii* predicted proteome in ToxoDB[6] and by profile-based homology searches using HHpred[5].

**Supplementary Table 2. Coq3 and Coq5 homologous sequences used for the phylogenetic analyses shown in Fig. 1E and Fig. 1F.**

| Phylogeny | Organism | Genebank/VEuPathDB | Reference |
|---|---|---|---|
| *Coq3* | *Toxoplasma gondii* | TGME49_266850 | This work |
| | *Hammondia hammondi* | HHA_266850 | Predicted |
| | *Neospora caninum* | NCLIV_039200 | Hypothetical |
| | *Besnoitia besnoiti* | BESB_049710 | Predicted |
| | *Cystoisospora suis* | CSUI_009995 | Predicted |
| | *Sarcocystis neurona* | SN3_01400045 | Predicted |
| | *Eimeria tenella* | ETH2_1576500 | Predicted |
| | *Cyclospora cayetanensis* | cyc_08631 | Hypothetical |
| | *Plasmodium falciparum* | PF3D7_0724300 | Predicted |
| | *Plasmodium cynomolgi* | PCYB_032810 | Predicted |
| | *Cryptosporidium hominis* | Chro.20298 TU502 | Predicted |
| | *Trypanosoma cruzi* | TcCLB.506147.60 | Predicted |
| | *Leishmania donovani* | LdBPK_354320.1 | Predicted |
| | *Aspergillus campestris* | XP_024691259.1 | Predicted |
| | *Candida albicans* | C6_01840C_A | Predicted |
| | *Saccharomyces cerevisiae* | P27680 | (7) |
| | *Rhizophagus irregularis* | RhiirFUN_008012 | Predicted |
| | *Homo sapiens* | ENSG00000132423 | (8) |
| | *Bos taurus* | Q3T131 | Predicted |
| *Coq5* | *Toxoplasma gondii* | TGME49_295690 | This work |
| | *Hammondia hammondi* | HHA_295690 | Predicted |
| | *Neospora caninum* | NCLIV_002130 | Predicted |
| | *Besnoitia besnoiti* | BESB_026650 | Predicted |
| | *Cystoisospora suis* | CSUI_003480 | Predicted |
| | *Cyclospora cayetanensis* | cyc_06773 | Predicted |
| | *Eimeria tenella* | ETH2_1538600 | Predicted |
| | *Cryptosporidium hominis* | TU502_Chro.10320 | Hypothetical |
| | *Plasmodium falciparum* | PF3D7_0204900 | Predicted |
| | *Mus musculus* | NP_080780.1 | Predicted |
| | *Homo sapiens* | ACD75052.1 | (9) |
| | *Saccharomyces cerevisiae* | AJS82729.1 | (10) |
| | *Candida albicans* | C2_05470W_A | Predicted |
| | *Trypanosoma cruzi* | TcCLB.510285.60 | Predicted |
| | *Trypanosoma brucei* | Tb11.01.2280 | Predicted |
| | *Leishmania donovani* | LdBPK_365590.1 | Predicted |
| | *Leishmania major* | LmjF.36.5360 | Predicted |

| | | | |
|---|---|---|---|
| UbiE | *E. coli* | KP670837.1 | (11) |
| UbiG | *E. coli* | NP_416735.1 | (12) |
| COMT | *Homo sapiens* | NG_011526.1 | (13) |

**Supplementary Table 3. Proteins identified from TgCoq5-TID mass spectrometry**

The excel sheet is in a separate document due to its large size. The excel file includes all the peptides detected in TgCoq5-TID and FbxO14-TID streptavidin IP (conditions and analysis included in the methods section) with their respective geneID, number of hits per sample, fold change and p-values.

The dataset was uploaded to PRIDE: **PXD059833**

**Supplementary Table 4. GenBank and EuPathDB sequence identifiers used for the phylogenetic analysis of Coq6, Coq7, and plant and bacterial ubiquinone synthesis monooxygenases shown in Fig. 5.**

| Organism | Genebank/EuPathDB | Reference |
|---|---|---|
| *Toxoplasma gondii* | TGME49_232540 | This work |
| *Plasmodium falciparum* | PF3D7_0815300 | FAD-dependent monooxygenase, putative |
| *Cryptosporidium hominis* | Chro.80315 | hypothetical |
| *Eimeria tenella* | ETH2_1101300 | FAD-dependent monooxygenase, putative |
| *Hammondia hammondi* | HHA_232540 | FAD binding domain-containing protein |
| *Neospora caninum* | NCLIV_032510 | FAD binding domain-containing protein |
| *Chromera velia* | Cvel_13198 | Tetracenomycin polyketide synthesis hydroxylase, putative |
| *Homo sapiens* | ENSG00000119723 | (14) |
| *Saccharomyces cerevisiae* | YGR255C | (15, 16) |
| *Arabidopsis thaliana* | At1g24340 | (17) |
| *Arabidopsis thaliana* | At3g24200 | (18) |
| *Escherichia coli* | P75728 | (19) |
| *Escherichia coli* | P25534 | (20) |
| *Escherichia coli* | P25535 | (21) |
| *Homo sapiens* | Q99807 | (22) |
| *Saccharomyces cerevisiae* | P41735 | (23) |
| *Oryza sativa Japonica group* | XP_015620261.1 | Uncharacterized protein |
| *Oryza sativa Japonica group* | XP_015632146. | 1 ubiquinone biosynthesis monooxygenase COQ6 |
| *Ostreococcus tauri* | XP_003079408.2 | Monooxygenase, FAD-binding |
| *Ostreococcus tauri* | XP_003081515.1 | Ubiquinone biosynthesis hydroxylase,UbiH/UbiF/VisC/COQ6 |
| *Chlorella variabilis* | XP_005850321.1 | hypothetical protein CoqF |
| *Chlorella variabilis* | XP_005852013.1 | hypothetical protein Coq6 |
| *Drosophila melanogaster* | NP_651967.2 | coenzyme Q7 |
| *Drosophila melanogaster* | NP_608934.1 | coenzyme Q6 |
| *Aspergillus bombycis* | XP_022384928.1 | ubiquinone biosynthesis protein Coq7 |
| *Aspergillus bombycis* | XP_022390693.1 | putative ubiquinone biosynthesis monooxgenase (Coq6) |

**Supplementary Table 5. Genebank sequence identifiers used for the phylogenetic tree in Fig S8.**

| Organism | Genebank/EuPathDB | Annotation |
|---|---|---|
| *Hyaloraphidium curvatum* | KAI9034245.1 | FAD binding domain-containing protein |
| *Ectocarpus siliculosus* | CBJ25915.1 | Monooxygenase |
| *Scytosiphon promiscuus* | CAM9992232.1 | unnamed protein product |
| *Oryza meyeriana var. granulata* | KAF0919747.1 | hypothetical protein |
| *Citrus sinensis* | KAH9772730.1 | FAD binding 3 domain-containing protein |
| *Desulfobacterales bacterium* | MDJ0831795.1 | FAD-dependent monooxygenase |
| *Acanthamoeba castellanii* | ACA1_006220 | FAD dependent monooxygenase |
| *Aphanomyces euteiches* | AeMF1_016247 | unspecified product |
| *Phytophthora capsici* | DVH05_022233 | unspecified product |
| *Globisporangium irregulare* | PIR_ G008400 | Polyketide hydroxylase |
| *Alloalcanivorax gelatiniphagus* | WP/9-172 205717242.1 | FAD-dependent monooxygenase |
| *Kwoniella newhampshirensis* | XP/4-325 066800455.1 | hypothetical protein |
| *Candidatus Nitrosotalea sp.* | HUM16632.1/6-383 | FAD-dependent monooxygenase |
| *Dendronephthya gigantea* | XP/6-385 028410004.1 | FAD-dependent monooxygenase apdD-like |
| *Paramuricea clavata* | CAB3997534.1/7-387 | FAD-monooxygenase |
| *Ramlibacter sp.* | MEO5670966.1 | FAD-dependent monooxygenase |
| *Betaproteobacteria bacterium* | MEI6468365.1 | FAD-dependent monooxygenase |
| *Heterostelium album PN500* | XP/19-415 020429748.1 | hypothetical protein |
| *Adineta vaga* | ACD54778.1 | FAD-binding monooxygenase-like protein |
| *Cyanidium caldarium* | KAK4534342.1 | hypothetical protein |
| *Xanthobacteraceae bacterium* | MDR3468332.1 | FAD-dependent monooxygenase |
| *Aspergillus lucknowensis* | XP/5-386 070890239.1 | FAD binding domain-containing protein |
| *Thetidibacter halocola* | WP/389-770 249200398.1 | FAD-dependent monooxygenase |
| *Acinetobacter seifertii* | WP/6-153 228271463.1 | FAD-dependent monooxygenase |
| *Dictyostelium discoideum AX4* | XP/30-399 636994.1 | FAD-binding monooxygenase |
| *Candidatus Methanoperedens nitratireducens* | WP/5-289 052368495.1 | FAD-dependent monooxygenase |
| *Haladaptatus pallidirubidus* | WP/7-215 390184515.1 | FAD-dependent monooxygenas |
| *Toxoplasma gondii* | TGG1_220360 | FAD-binding domain containing protein |
| *Linnemannia gamsii* | KAK3838246.1 | hypothetical protein |
| *Planctomyces sp.* | WP/5-353 075094984.1 | FAD-dependent monooxygenase |
| *Cyanobacteria bacterium* | MDB5100591.1 | hypothetical protein |
| *Deltaproteobacteria bacterium* | TMA90433.1 | FAD-dependent oxidoreductase |

### Supplementary Table 6. Proteins identified from TgCoqFAD-3HA mass spectrometry

The excel sheet is in a separate document due to its large size. The excel file includes all the peptides detected in TgCoqFAD-3HA and TgCoq1-3HA IP (conditions and analysis included in the methods section) with their respective geneID, number of hits per sample, fold change and p-values.

The dataset was uploaded to PRIDE: PXD071065.

**Supplementary Table 7. Primers used in this study**

| Cell line name | Primer name | sequence |
|---|---|---|
| *TgCoq3-3HA* | 3' CRISPR F | GcgtcttgtcccgccttcagGTTTTAGAGCTAGAAATAGCAAG |
| | HA homology F | CTTTTGCAGGTTCTCTTTTGCAATGGCATTTGAGAAAATCTACCCGTACGACGTCCCGGA |
| | HA homology R | acatcctgcgtgctgctgtatgcttgacgcgaaaatgtgtCCCTCGGGGGGGCAAGAATT |
| *TgCoq5-3HA* | 3' CRISPR F | cggaatgtgttactgagttgGTTTTAGAGCTAGAAATAGCAAG |
| | HA homology F | GACCGTTGCCATACACTCAGGGTTCAAGTTAGGAAACCTGTACCCGTACGACGTCCCGGA |
| | HA homology R | tccgcgttctccgaatgttgcagaactgtccttgttacccCCCTCGGGGGGGCAAGAATT |
| *iΔ3Ty-TgCoq3* | 5' CRISPR F | ttgccagcaacctcacgcgtGTTTTAGAGCTAGAAATAGCAAG |
| | PI ty homology F | tctagagcggtgtcaatgctccgaaaccggttgatatgtaaagcttcgccaggctgtaaatc |
| | PI ty homology R | CCTTACAGCAAGTTGAAGAAAAGGTTGAAGGTGTCTTCATGTCCAGGGGATCCTGATTG |
| *iΔTgCoq3* | PI no tag homology F | tctagagcggtgtcaatgctccgaaaccggttgatatgtaaagcttcgccaggctgtaaatc |
| | PI no tag homology R | CCTTACAGCAAGTTGAAGAAAAGGTTGAAGGTGTCTTCATggttgaagacagacgaaagc |
| *iΔTgCoq5* | 5' CRISPR F | ctcagttgtcgtgccaagctGTTTTAGAGCTAGAAATAGCAAG |
| | PI no tag homology F | ttctgttcgtttttgctcatccgttcaagctaccaggaagcttcgccaggctgtaaatc |
| | PI homology R | ATGAGATCTTTCGCGGTTCCTTGTGCGCGTTGCGGGGGGGggttgaagacagacgaaagc |
| *TgCoq3-TID* | HA homology F | CTTTTGCAGGTTCTCTTTTGCAATGGCATTTGAGAAAATCTACCCGTACGACGTCCCGGA |
| | HA homology R | acatcctgcgtgctgctgtatgcttgacgcgaaaatgtgtCCCTCGGGGGGGCAAGAATT |
| *TgCoq5-TID* | HA homology F | GACCGTTGCCATACACTCAGGGTTCAAGTTAGGAAACCTGTACCCGTACGACGTCCCGGA |
| | HA homology R | tccgcgttctccgaatgttgcagaactgtccttgttacccCCCTCGGGGGGGCAAGAATT |
| *TgCoq4-3HA* | 3' CRISPR F | /5Phos/AAGTTGCACTCTCTGCGCTTCAGTCCG |
| | 3' CRISPR R | /5Phos/AAAACGGACTGAAGCGCAGAGAGTGCA |
| | HA homology F | TGCCCACCTCGCGCCTCACGTAGCTCGCGACCAAAGTTCTTACCCGTACGACGTCCCGGA |
| | HA homology R | TCTTCCGGAAAAGTGAAGAGTGTGTCGAAACAAAGAGAGACCCTCGGGGGGGCAAGAATT |
| *iΔTgCoq4-3HA* | 5' CRISPR F | /5Phos/AAGTTGTACTCCACACAGACGGGGGG |
| | 5' CRISPR R | /5Phos/AAAACCCCCCGTCTGTGTGGAGTACA |
| | PI homology F | AGCGGCAGATCCTTGCGTGGAGAGGCGTCGCTCACTGCTGaagcttcgccaggctgtaaa |
| | PI homology R | GAGAGAGGCGCAGAAGGCGCCGAAGAATGGGTCGCCTCATggttgaagacagacgaaagc |
| *TgCoq6-3HA* | 3' CRISPR F | GATCTATTGTGCTTCTGGTGGTTTTAGAGCTAGAAATAGCAAG |
| | HA homology F | TGTGGAAGCCGTTGTAAAAGATATTCTACAGAGGCTCGCTTACCCGTACGACGTCCCGGA |

| | | |
|---|---|---|
| | HA homology R | TTATCCCCAGAAGAAGACCGTCGTGCCACCGGCTCTGAGTCCCTCGGGGGGGCAAGAATT |
| *iΔTgCoq6-3HA* | 5' CRISPR F | /5Phos/AAGTTGTCGTCACCCACTCTTTCCCG |
| | 5' CRISPR R | /5Phos/AAAACGGGAAAGAGTGGGTGACGACA |
| | PI homology F | TTCCCGTGCAGTTTCGATTTAAAGCCATTCCTAAGGCTCTaagcttcgccaggctgtaaa |
| | PI homology R | CTGTGGCACCGTGGCGCCTTCCAGCTAAAAACGGTGCCATggttgaagacagacgaaagc |
| *TGGT1_236140-3HA* | 3' CRISPR F | /5Phos/AAGTTGAGCAGCGGTGTGTACTCGCG |
| | 3' CRISPR R | /5Phos/AAAACGCGAGTACACACCGCTGCTCA |
| | HA homology F | GCGACACCAGCTGGCGCTCACCAGACCTCCAGCTGAAGAATACCCGTACGACGTCCCGGA |
| | HA homology R | CGAAACTGTTTCGACACAGTGCTGCGCCTTTCAGAGGTTCCCCTCGGGGGGGCAAGAATT |
| *TGGT1_270150-3HA* | 3' CRISPR F | /5Phos/AAGTTGACATCACGAGAGTGAAGAGG |
| | 3' CRISPR R | /5Phos/AAAACCTCTTCACTCTCGTGATGTCA |
| | HA homology F | GGAGAGAGACGAGTTTCTTCGAAGTCTCACAAATCTCTCGTACCCGTACGACGTCCCGGA |
| | HA homology R | AAGGCCGCGACGCCTTTTTCTCTGCGTGTGCCCGCGCTTTCCCTCGGGGGGGCAAGAATT |
| *TGGT1_236510-3HA* | 3' CRISPR F | /5Phos/AAGTTGTAGCTAGGTGTATGTACATG |
| | 3' CRISPR R | /5Phos/AAAACATGTACATACACCTAGCTACA |
| | HA homology F | CAAGAGGAGGCTCGAACATCAAACTGACGCTTCGCTCTACTACCCGTACGACGTCCCGGA |
| | HA homology R | ATCGACAAAACGCTGTATATGCCTTTGCATGATGAGGATTCCCTCGGGGGGGCAAGAATT |
| *TGGT1_239610-3HA* | 3' CRISPR F | /5Phos/AAGTTGACAAGCTGTAGACTGGCTTTG |
| | 3' CRISPR R | /5Phos/AAAACAAAGCCAGTCTACAGCTTGTCA |
| | HA homology F | CGTCGTTCTGCGGAGACGCCTCAGACTTATTCCTGGAATCTACCCGTACGACGTCCCGGA |
| | HA homology R | TGGGATAGCCGGCCGGTCACTCTCCCCGCGCTCCCAGCTCCCCTCGGGGGGGCAAGAATT |
| *TGGT1_288710-3HA* | 3' CRISPR F | /5Phos/AAGTTGAGCGTATAGAAGAGTGAGCG |
| | 3' CRISPR R | /5Phos/AAAACGCTCACTCTTCTATACGCTCA |
| | HA homology F | GAGAGAAAAACTTCGAAGTCTCGGCGTTCAGCAAAGCCACTACCCGTACGACGTCCCGGA |
| | HA homology R | GCATGTGTATAAAAATAAATACATACACACATGCACAGAGCCCTCGGGGGGGCAAGAATT |
| *TGGT1_263840-3HA* | 3' CRISPR F | GGTGGAGTGTGTGTGGCGATGTTTTAGAGCTAGAAATAGC |
| | HA homology F | GACTGGACCCGAAGAATGGGCATGGTCAGATCCTACTCGATACCCGTACGACGTCCCGGA |
| | HA homology R | ACGGCTCAGGCAAAGACACCCATCCAGCCCGCACTGCCTTCCCTCGGGGGGGCAAGAATT |
| *TGGT1_251890-3HA* | 3' CRISPR F | ACAGATACATGTGAACTCACGTTTTAGAGCTAGAAATAGC |
| | HA homology F | AGAGTCACCAGCGGGTACGACACTCCTGCGTGTCAAGATCTACCCGTACGACGTCCCGGA |
| | HA homology R | TGACACGGTGCTTGCTTTTAAGATATACCTGCGCAAGTCCCCTCGGGGGGGCAAGAATT |

| | | |
|---|---|---|
| *iΔTGGT1_236510-3HA* | 5' CRISPR F | /5Phos/AAGTTGAACAGCCATGCGTCGCTGCTG |
| | 5' CRISPR R | /5Phos/AAAACAGCAGCGACGCATGGCTGTTCA |
| | PI homology F | TGTATACGTATATATGTGTGTGTAAGAGGATGTACGAAGCaagcttcgccaggctgtaaa |
| | PI homology R | CACGGCTTGTCCTCCTCGCATATCCCACAGAGATCTGCATggttgaagacagacgaaagc |
| *iΔTGGT1_239610-3HA* | 5' CRISPR F | /5Phos/AAGTTGCTCACATGTTCTTCGTCTTG |
| | 5' CRISPR R | /5Phos/AAAACAAGACGAAGAACATGTGAGCA |
| | PI homology F | GTACATCCATATATGTATACATACATGTGCGTGTGTCAGAaagcttcgccaggctgtaaa |
| | PI homology R | ACGGCGACGCAAAGACGCGCAGTGAGCGCAGAGGCGCCATggttgaagacagacgaaagc |
| *iΔTGGT1_288710-3HA* | 5' CRISPR F | /5Phos/AAGTTGGGCTTTCCCAGCCGACTCAG |
| | 5' CRISPR R | /5Phos/AAAACTGAGTCGGCTGGGAAAGCCCA |
| | PI homology F | TTCACCAACCGCTTCTCTCTTTGGTGGCTGCGCGCCCGCGaagcttcgccaggctgtaaa |
| | PI homology R | CTCTCCGTCGCGGCGGGTGGAAGAGCCGCAGCGCCACCATggttgaagacagacgaaagc |
| *iΔTGGT1_251890-3HA* | 5' CRISPR F | /5Phos/AAGTTGCTTGGTCGACAAGCAAAAGG |
| | 5' CRISPR R | /5Phos/AAAACCTTTTGCTTGTCGACCAAGCA |
| | PI homology F | GAAGCAACTCACTTTCTCTGCAATGACAGTCGCGTTCCTAaagcttcgccaggctgtaaa |
| | PI homology R | GGAACGCCAAAGGACGGGGGAAAAGTTGTGTGAAGGCCATggttgaagacagacgaaagc |
| pQE80L_TgCoq3 | Coq3-BamHI-F | CGCGGATCCGCGATGAAGACACCTTCAACCTTTTCTTCAAC |
| | Coq3-KpnI-R | CGGGGTACCCCGTCAGATTTTCTCAAATGCCATTGC |
| pQE80L_TgCoq5 | Coq5 BamHI F | CACGGATCCGCGATGAGATCTTTCGCGGTTCCTTG |
| | Coq5 KpnI R | GGGGGTACCCTACAGGTTTCCTAACTTGAACCCTGAGTGT |
| CRISPR sequencing | M13 R | CAGGAAACAGCTATGAC |
| | pSS013/CRISPR R | GGTCCAACCAGGATCACCAAC |
| Complementation plasmid | pK016 F | ATCAGGATCCCCTGGACTAAatcaccgttgtgctcacttc |
| | pK016 R | CCAGCTAAAAACGGTGCCATggttgaagacagacgaaagc |
| | CoqFAD F | gctttcgtctgtcttcaaccATGGCACCGTTTTTAGCTGG |
| | CoqFAD R | tcctgGTTGGTGTGCACCTCAGCGAGCCTCTGTAGAATAT |
| | Ty F | ATATTCTACAGAGGCTCGCTGAGGTGCACACCAACcagga |
| pcAtFMO | pc_AtF_F | CGATCATCCTGGGGAAGCAGGAGGTGCACACCAACcagga |
| | pc_AtF_R | ACAGGGAGCTTTGCCGCGTCCATCCTAGCAATCCAGGGAAAG |
| | AtF_F | TTCCCTGGATTGCTAGGATGGACGCGGCAAAGCTCCCTGT |
| | AtF_R | tcctgGTTGGTGTGCACCTCCTGCTTCCCCAGGATGATCGT |
| pcAtCoq6 | pc_At6_F | AGTCACTGCCCCTCTTTTCAGAGGTGCACACCAACcagga |
| | pc_At6_R | GCAATGTCGTGCTGAGGCCCCCTAGCAATCCAGGGAAAGAG |

| | | |
|---|---|---|
| | At6_F | TCTTTCCCTGGATTGCTAGGGGGCCTCAGCACGACATTGC |
| | At6_R | tcctgGTTGGTGTGCACCTCTGAAAAGAGGGGCAGTGACTG |
| | Ty R | gaagtgagcacaacggtgatTTAGTCCAGGGGATCCTGATT |
| UPRT complementation repair | SAG4 UPRT F | gcctgtacgccggttgtcgcggtcgtttgcagattgctttacgccgctgagactaactag |
| | SAG4 UPRT R | cgtcgttcttcgcgagggttacagcaccgattcgaagaaaccctcgggggggcaagaatt |
